# Circuits that encode and predict alcohol associated preference

**DOI:** 10.1101/578401

**Authors:** Kristin M. Scaplen, Mustafa Talay, Sarah Salamon, Kavin M. Nuñez, Amanda G. Waterman, Sydney Gang, Sophia L. Song, Gilad Barnea, Karla R. Kaun

## Abstract

Substance use disorders are chronic relapsing disorders often impelled by enduring memories and persistent cravings. Alcohol, as well as other addictive substances, remolds neural circuits important for memory to establish obstinate preference despite aversive consequences. How pertinent circuits are selected and shaped to result in these unchanging, inflexible memories is unclear. Using neurogenetic tools available in *Drosophila melanogaster* we define how circuits required for alcohol associated preference shift from population level dopaminergic activation to select dopamine neurons that predict behavioral choice. During memory expression, these dopamine neurons directly, and indirectly via the mushroom body (MB), modulate the activity of interconnected glutamatergic and cholinergic output neurons. Transsynaptic tracing of these output neurons revealed at least two regions of convergence: 1) a center of memory consolidation within the MB implicated in arousal, and 2) a structure outside the MB implicated in integration of naïve and learned responses. These findings provide a circuit framework through which dopamine neuron activation shifts from reward delivery to cue onset, and provides insight into the inflexible, maladaptive nature of alcohol associated memories.

## Introduction

An organism’s behavior is guided by memories of past experiences and their associated positive or negative outcomes. The accuracy of these associations is vital to the organism’s survival. In a changing environment, however, successful associations should be dynamic and malleable, providing opportunities for updating associations based on new information. In substance use disorders (SUD) this flexibility is often absent or difficult to achieve (Font and Cunningham 2012, Torregrossa and Taylor 2013). Preference and cravings for drugs of abuse, including alcohol, persist in the face of aversive consequences leading to maladaptive drug seeking behaviors and a devastating economic and social impact on individuals, communities, and society as a whole. Understanding the circuitry mechanisms that underlie how these memories are encoded and retrieved is critical to understanding why these memories are so resistant to change.

Systems memory consolidation suggests that individual brain regions have a time limited role in the recall of memories as they are slowly reorganized and transferred to anatomically distinct regions in the brain. More recent work has suggested that even within a brain region, such as the hippocampal formation, circuits important for encoding and retrieval are distinct (Roy, Kitamura et al. 2017, Cembrowski, Phillips et al. 2018). Strikingly, drugs of abuse, such as alcohol disrupt these memory systems resulting in enduring preferences, attentional bias for associated cues (Field and Cox 2008, Fadardi, Cox et al. 2016), and habitual behaviors. The neuronal genetic, morphologic, and physiologic diversity that exists within the brain, however, has precluded mammalian animal models from accessing the level of spatial resolution in a temporally specific manner required to identify and investigate microcircuits important for alcohol associated behaviors. We sought to address how drugs of abuse can shift memory from being flexible to inflexible by using a combination of behavioral, thermogenetic, *in vivo* calcium imaging, and transsynaptic tracing techniques in *Drosophila melanogaster.*

The neural circuitry underlying the *Drosophila* reward response is complex and remarkably similar to mammals (Scaplen and Kaun 2016). *Drosophila* exhibit sophisticated behaviors in response to changes in their environment, including formation of appetitive memories for food associated with environmental cues (Waddell 2010, Kahsai and Zars 2011, Schleyer, Saumweber et al. 2011, Herrero 2012, Fernandez and Kravitz 2013, Heisenberg 2015, Owald, Lin et al. 2015). *Drosophila* also form appetitive memories for the pharmacological properties of alcohol (Kaun, Azanchi et al. 2011, Nuñez, Azanchi et al. 2018). These alcohol memories are persistent and impel flies to walk over a 120V electric shock in the presence of associated cues. This type of goal directed behavior, in the face of aversive consequences, suggest that alcohol associated memories are more inflexible than memories for natural reward like sucrose (Kaun et al, 2011). *Drosophila* provides an ideal model to investigate these differences as the neurogenetic tools available in this species permits dissection of memory circuits with unprecedented temporal and spatial resolution.

In *Drosophila* the establishment of alcohol preference requires an associative central brain structure called the mushroom bodies (MB) and dopamine (DA) (Kaun, Azanchi et al. 2011). We show that circuits required for formation of alcohol preference shift from population-level dopaminergic encoding to two microcircuits comprising of interconnected dopaminergic, glutamatergic, and cholinergic neurons. These circuits converge onto the fan-shaped body (FSB), a higher-order brain center implicated in arousal and modulating behavioral response (Liu, Seiler et al. 2006, Weir, Schnell et al. 2014, Weir and Dickinson 2015, Pimentel, Donlea et al. 2016, Qian, Cao et al. 2017, Donlea, Pimentel et al. 2018, Hu, Peng et al. 2018, Troup, Yap et al. 2018). Our results, therefore, provide an *in vivo* circuit framework for how drugs of abuse temporally regulate acquisition and expression of sensory memories, which ultimately results in a shift in behavioral response from malleable to inflexible.

## Results

### Dopamine neurons innervating the mushroom body are required for alcohol reward associations

Dopamine has a long-standing role in addiction and a defined role in reward-related behavioral learning that spans across species (Yoshimoto, McBride et al. 1992, Robbins and Everitt 2002, Hyman, Malenka et al. 2006, Wanat, Willuhn et al. 2009, Kaun, Azanchi et al. 2011, Torregrossa, Corlett et al. 2011, Scaplen and Kaun 2016). In mammals, midbrain ventral tegmental area (VTA) and substantia nigra pars compacta (SNc) structures are primary sources of dopamine responsible for modulating synaptic plasticity within the brain. However, the neuronal heterogeneity that exists within these structures, and lack of necessary spatial and temporal resolution to parse apart subpopulations or isolate individual neurons in behaving animals, have precluded the field from gaining a complete understanding of how dopaminergic neurons engage and augment underlying neural activity both in normal learning and addiction.

We first sought to identify which dopaminergic neurons within the *Drosophila* brain are necessary for alcohol associated preference. In *Drosophila* a discrete population of protocerebral anterior medial (PAM) dopamine neurons have an identified role in detecting and processing natural rewards. PAM neurons are required for the acquisition of sucrose memory (Liu, Placais et al. 2012, Huetteroth, Perisse et al. 2015, Yamagata, Ichinose et al. 2015), and known to be activated by sucrose administration (Liu, Placais et al. 2012, Harris, Kallman et al. 2015). We first tested the requirement of activity of PAM dopamine neurons in alcohol associative preference (Figure 1a). For selective manipulations of these neurons, we expressed the dominant negative temperature sensitive *shibire* (*shi*^*ts1*^) using R58E02-GAL4 (Figure 1b). To establish temporal requirements, we temporarily and reversibly inactivated neurotransmission by raising the temperature to restricted levels (30°C) during acquisition, the overnight consolidation period, or retrieval (Figure 1a). Acquisition was defined as the time during which an odor was presented in isolation (unpaired odor) and a second odor was paired with an intoxicating dose of vaporized ethanol (paired odor). During acquisition, reciprocally trained flies received three of these spaced cue (odor) and overlapping cue sessions (odor + ethanol). Post-acquisition, flies were given a choice 24 hours later between the unpaired and paired odor in a Y maze (Figure 1a). Retrieval was defined as the time during which the flies chose between the unpaired and paired odors 24 hours post acquisition.

**Fig. 1.**
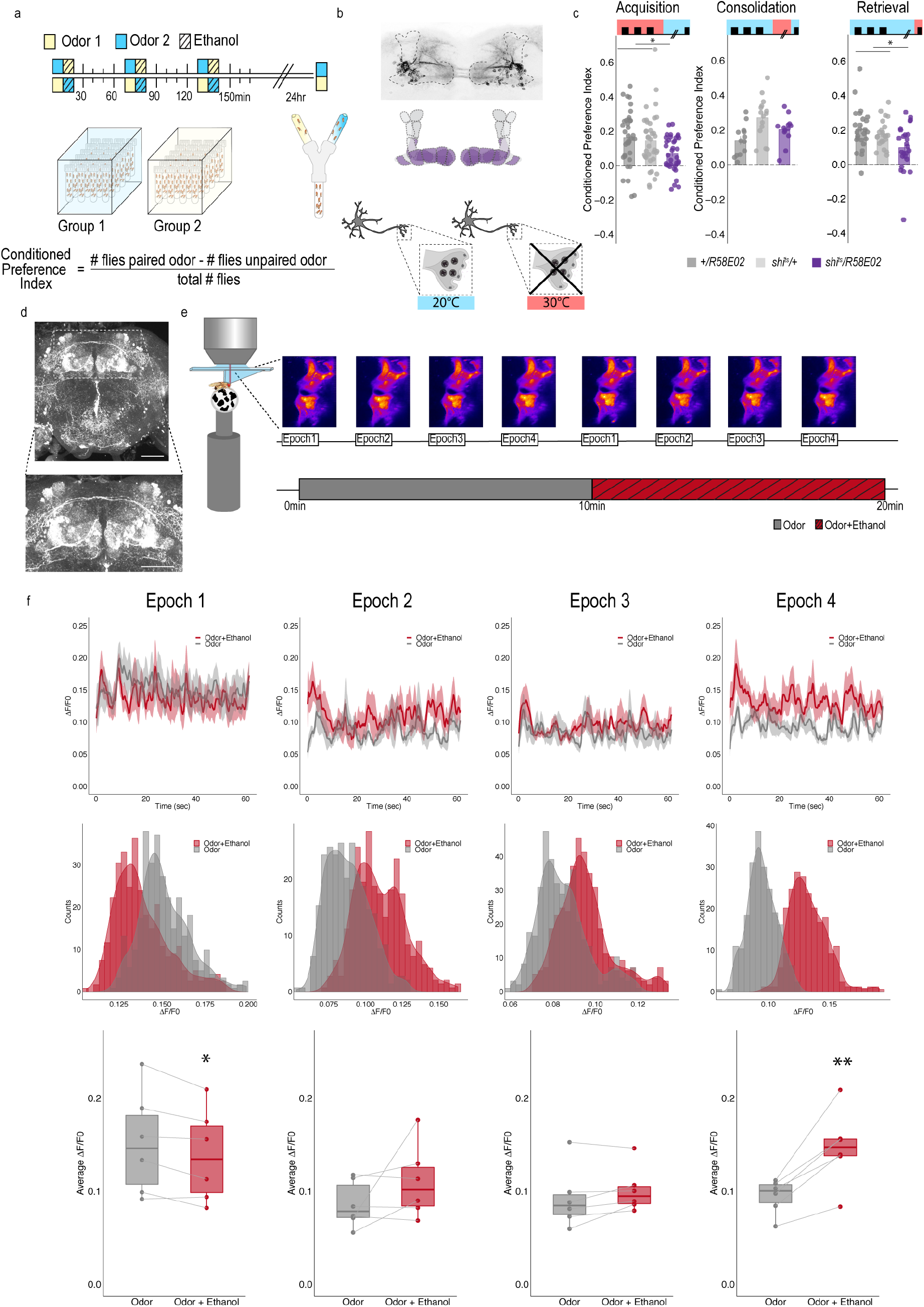
PAM dopaminergic neurons are necessary for encoding alcohol associated preference. (a) Schematic illustrating odor condition preference paradigm. Vials of 30 flies are presented with three sessions of 10 minutes of an unpaired odor, followed by 10 minutes of a paired odor plus intoxicating vaporized ethanol. To control for odor identity, reciprocal controls were used. (b) PAM dopaminergic neurons innervate the horizontal lobe of MB. Confocal stack illustrates innervation pattern using R58E02>MCD-GFP. Dashed line outlines MB region. MB schematic illustrates MB compartments innervated by PAM neurons. Lower schematic highlights mechanisms of thermogenetics using shibire^ts1^. Temperature is used to inactivate neurotransmission at restrictive temperatures (30°C), but not at permissive temperatures (20°C). (c) PAM dopaminergic neurons activity is necessary during acquisition and retrieval, but not consolidation. Bar graphs illustrate mean +/- standard error of the mean. Raw data are overlaid on bar graphs. Each dot is an n of 1, which equals approximately 60 flies (30 per odor pairing). One-way ANOVA with Tukey Posthoc was used to compare mean and variance. *p<0.05 (d) Example confocal stack labeling dopamine within the brain. Dopamine labeling is concentrated around the MB. (e) Schematic of calcium imaging protocol. Flies are exposed to odor followed by odor plus intoxicating vaporized ethanol while resting or walking on a ball. We used the same odor for both conditions so we could better compare circuit dynamics in response to ethanol and control for odor identity. Fluorescence was captured for 61sec recording epochs that were equally spaced by 2 minutes. Example maximum intensity fluorescent stacks are shown for each epoch. (f) Average traces recorded during the odor and odor plus ethanol epochs. Middle panels illustrate the binned ΔF/F0 and highlights a change in calcium dynamics as a consequence of ethanol exposure. Lower panels illustrate the average ΔF/F0 for each fly in each condition at each epoch. Within Subject Repeated Measures ANOVA with a Bonferroni Posthoc was used to compare mean and variance across condition and time. Scale bar = 50 μm *p<0.05 **p<0.01

Previous work established that flies show preference for the cues associated with ethanol intoxication 24 hour later, regardless of odor identity (Kaun et al 2011). We found inactivating neurotransmission in PAM dopaminergic neurons during acquisition or retrieval, but not during the overnight consolidation, significantly reduced preference for cues associated with ethanol (Figure 1c). Strikingly, despite dopamine’s established role in modulating locomotor and motor responses (Romo and Schultz 1990, Lima and Miesenbock 2005, Schultz 2007, Dodson, Dreyer et al. 2016, Howe and Dombeck 2016, Syed, Grima et al. 2016, da Silva, Tecuapetla et al. 2018), inactivating PAM dopaminergic neurons did not affect ethanol induced activity (Supplementary Figure 1). Together, these results suggest that neurons required for encoding preference are not required for the locomotor response to the acute stimulatory properties of ethanol. Formation of alcohol memories, instead, occurs within the circuit framework defined by memories for natural reward.

### Dopaminergic encoding of alcohol memory acquisition occurs at the population level

To determine how alcohol influenced activity of PAM dopaminergic neurons to shift memories from malleable to inflexible, we first used a dopamine staining protocol to label dopamine within the brain following 10 minutes of air or alcohol. As expected there was a significant amount of dopamine labeled within the mushroom body and the majority of fluorescence was limited to the horizontal lobes (Figure 1d). We hypothesized that dopamine fluorescence would increase within the horizontal lobes of the mushroom body in response to alcohol, however, quantification of fluorescence did not provide significant differences (Supplementary Figure 2). We reasoned that dopamine staining likely could not distinguish between dopamine in the presynaptic terminals and dopamine in the synaptic cleft, and thus turned to 2-photon functional calcium imaging to monitor circuitry dynamics of PAM dopaminergic activity in the context of intoxicating alcohol. We used *R58E02-Gal4* to express *GCaMP6m* (Chen, Wardill et al. 2013) and recorded from the PAM presynaptic terminals at the MB while naïve flies were presented with 10 minutes of odor, followed by 10 minutes of odor plus intoxicating doses of alcohol (Figure 1e).

**Fig. 2.**
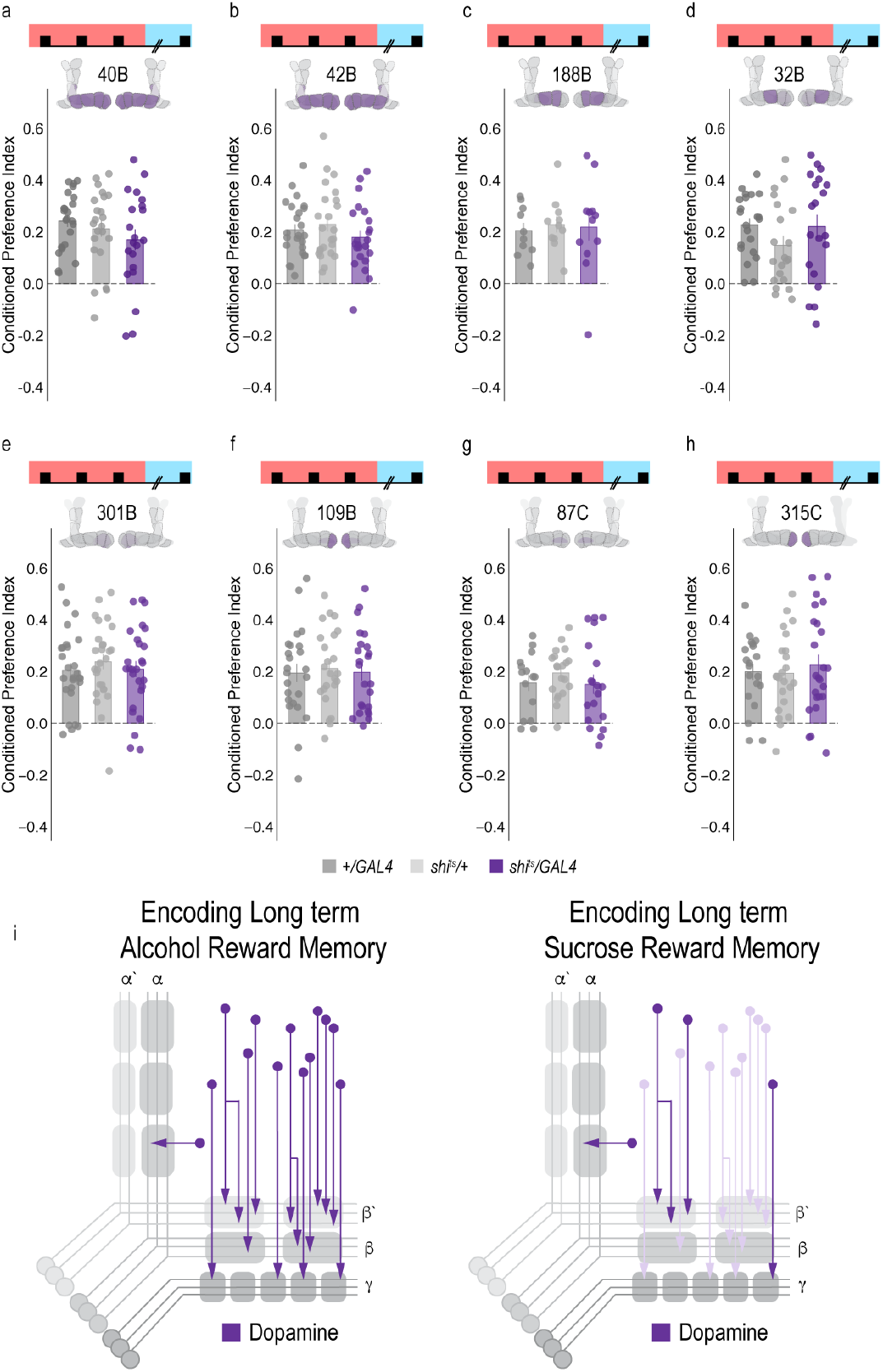
Subsets of PAM dopaminergic neurons are dispensable for encoding alcohol associated preference. **(a-h)** Using thermogenetics to inactivate neurotransmission during acquisition in dopaminergic neurons with varying expression patterns did not disrupt alcohol associated preference. Split-Gal4 lines tested are ordered by MB innervation patterns. Bar graphs illustrate mean +/- standard error of the mean. Raw data are overlaid on bar graphs. Each dot is an n of 1, which equals approximately 60 flies (30 per odor pairing). One-way ANOVA was used to compare mean and variance. (i) Model comparing encoding of alcohol reward memory and sucrose reward memory. Encoding alcohol reward memory requires population level PAM dopaminergic activity whereas sucrose reward memory is supported by subsets of PAM dopaminergic neurons during acquisition (Ichinose, Aso et al. 2015, Yamagata, Ichinose et al. 2015). The number of dopamine neurons recruited during acquisition may underlie why alcohol reward memories are persistent.

Interestingly, early in the respective recording sessions (odor vs odor + alcohol), changes in calcium dynamics was greater in the odor only group (Figure 1e, 1f), however with prolonged alcohol exposure, greater calcium dynamics started to emerge from the population of dopamine neurons in the odor + alcohol group (Figure 1f). This suggests that the pharmacological properties of alcohol increase dopaminergic neuronal activity in vivo, nicely complementing in vitro physiology in VTA slices that demonstrate VTA activity is increased by moderate alcohol doses (Brodie and Appel 1998, Morikawa and Morrisett 2010) and micro dialysis studies that report increased dopamine in the nucleus accumbens (Di Chiara and Imperato 1985, Imperato and Di Chiara 1986, Yoshimoto, McBride et al. 1992, Di Chiara, Tanda et al. 1996, Bassareo, De Luca et al. 2003, Howard, Schier et al. 2008), central amygdala (Yoshimoto, Ueda et al. 2000), and medial prefrontal cortex (Ding, Oster et al. 2011). However, the complexity of mammalian VTA circuits has obscured our understanding of whether increased dopamine release in downstream regions is due to more firing of discrete populations of VTA dopamine neurons, or recruitment of the larger population of VTA dopamine neurons.

To address whether specific subsets of dopamine neurons within the PAM neuron population are necessary for alcohol associated preference, we blocked transmission in subsets of these neurons using 18 highly specific split-Gal4 lines during both acquisition and retrieval. We found that preference was disrupted when neurotransmission was blocked in DA neurons projecting to the medial aspect of horizontal MB (Supplementary Figure 3). We therefore selected split-Gal4 lines that targeted the medial aspect of the horizontal lobe and determined their role specifically in acquisition of alcohol associated preference. Surprisingly, unlike long-term sucrose memory (Ichinose, Aso et al. 2015, Yamagata, Ichinose et al. 2015), thermogenetic inactivation of specific subsets of dopamine neurons innervating compartments of the medial horizontal lobe during acquisition did not disrupt 24-hour alcohol associated preference (Figure 2a-h, Supplementary Figure 4a). Cell counts of the broadest split-GAL4 lines (40B and 42B), HL9 and R58E02 driver lines revealed that despite targeting nearly all of the horizontal lobes of the MB, 40B, 42B, and HL9 expressed in significantly fewer cells (Supplementary Table 1). Together these data suggest that unlike long-term sucrose memory, subsets of dopamine neurons are not responsible for the acquisition of alcohol associated preference, and disruption of this process requires the disruption of population level dopaminergic activity. Thus, we speculate that alcohol memories are enduring and less adaptive than sucrose memories because alcohol engages a larger number of dopamine neurons during memory acquisition (Figure 2i).

**Table 1.**
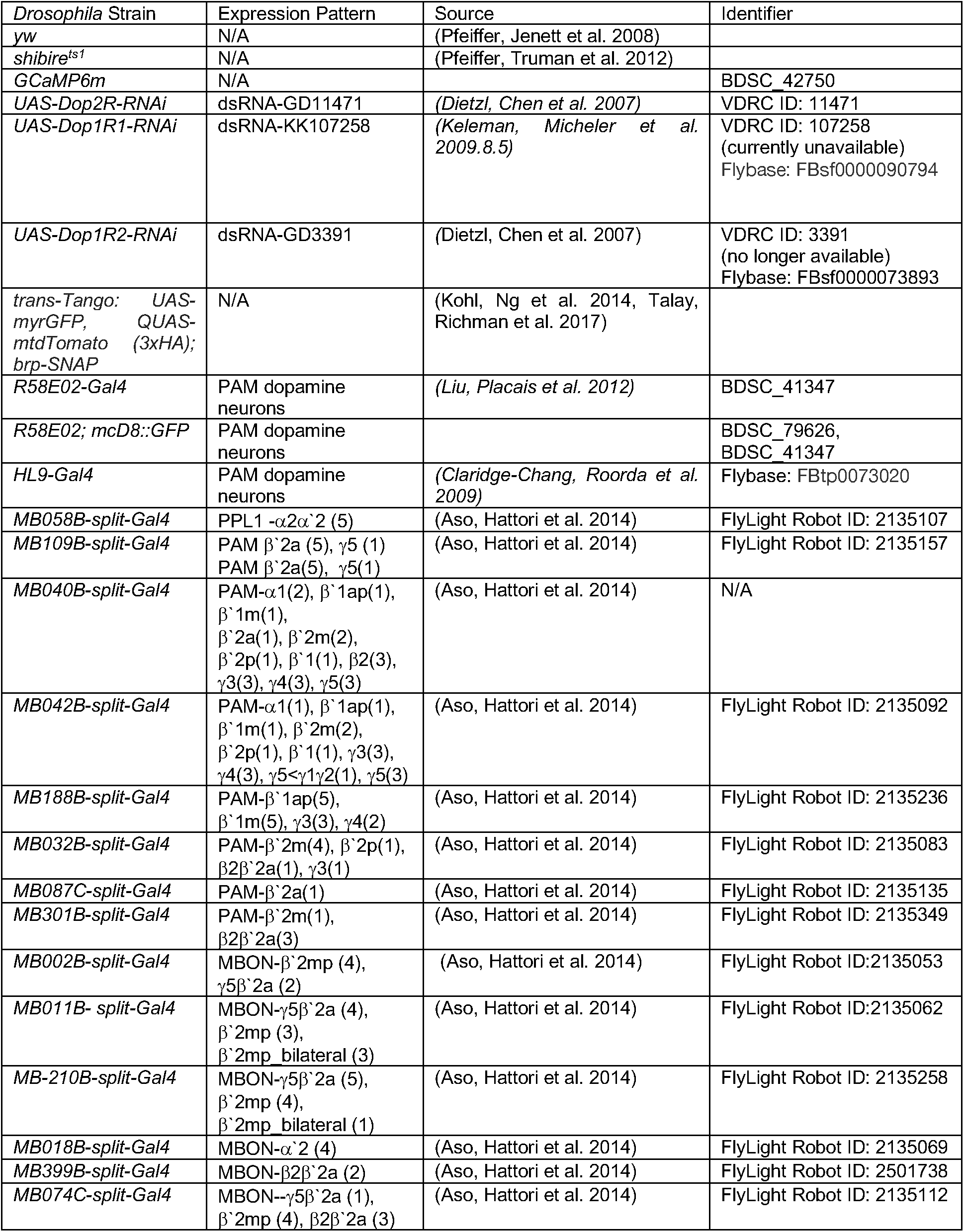
Fly Strains used in this manuscript.

**Fig. 3.**
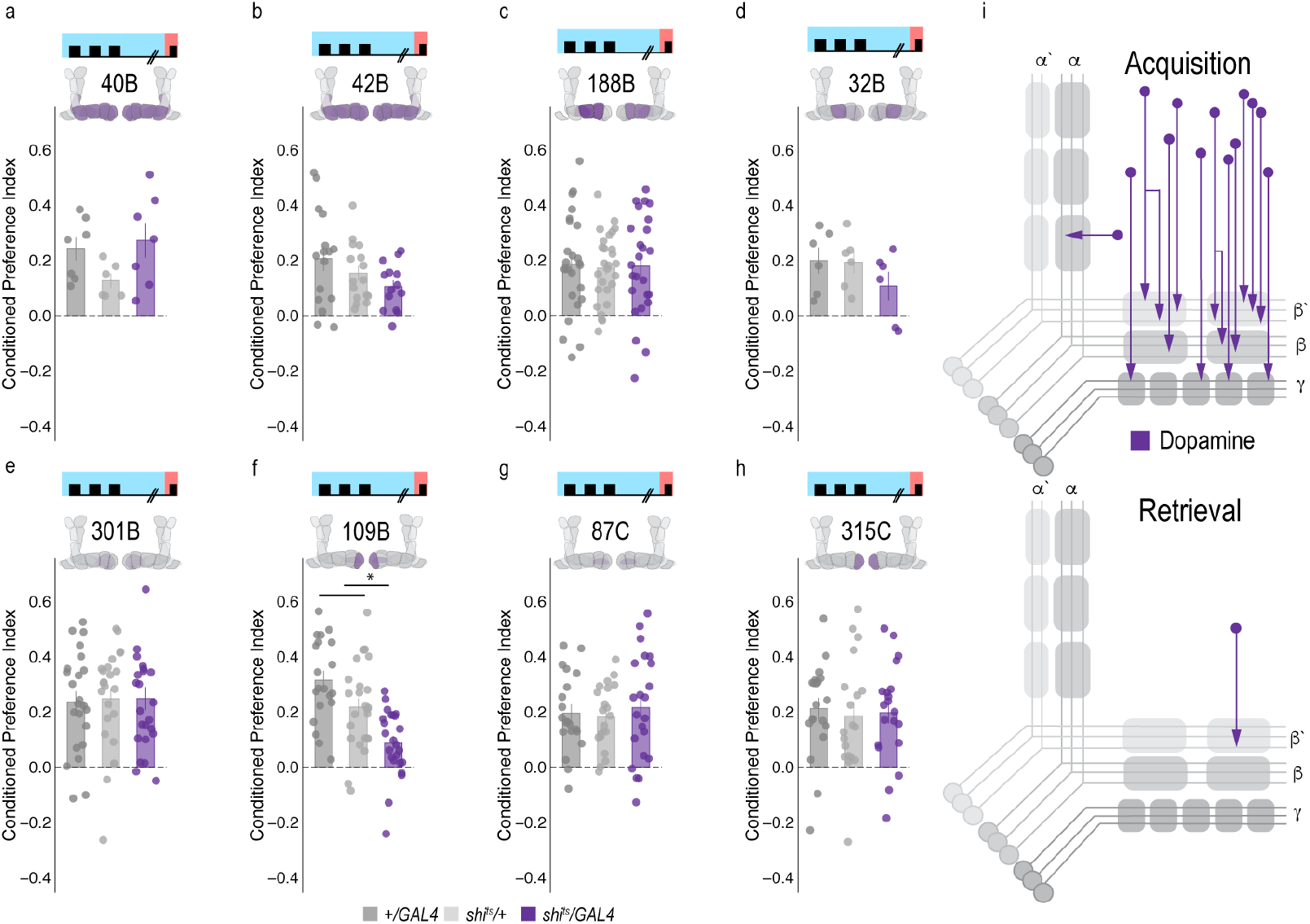
Memory expression during retrieval is dependent on a sparse population of dopamine neurons. (a-h) Thermogenetic were used to inactivate neurotransmission during retrieval, but not acquisition, in PAM dopaminergic neurons with varying expression patterns. (f) Inactivating β’2a dopamine neurons during retrieval significantly reduced preference for alcohol associated cues. Bar graphs illustrate mean +/- standard error of the mean. Raw data are overlaid on bar graphs. Each dot is an n of 1, which equals approximately 60 flies (30 per odor pairing). One-way ANOVA with Tukey Posthoc was used to compare mean and variance. *p<0.01 (i) Model of circuits responsible for encoding alcohol associated preference during acquisition and circuits responsible for the expression of alcohol associated preference during retrieval. This model highlights the importance of population level dopaminergic activity during acquisition, whereas sparse subsets of dopaminergic activity are important during retrieval for the expression of alcohol associated preference.

**Fig. 4.**
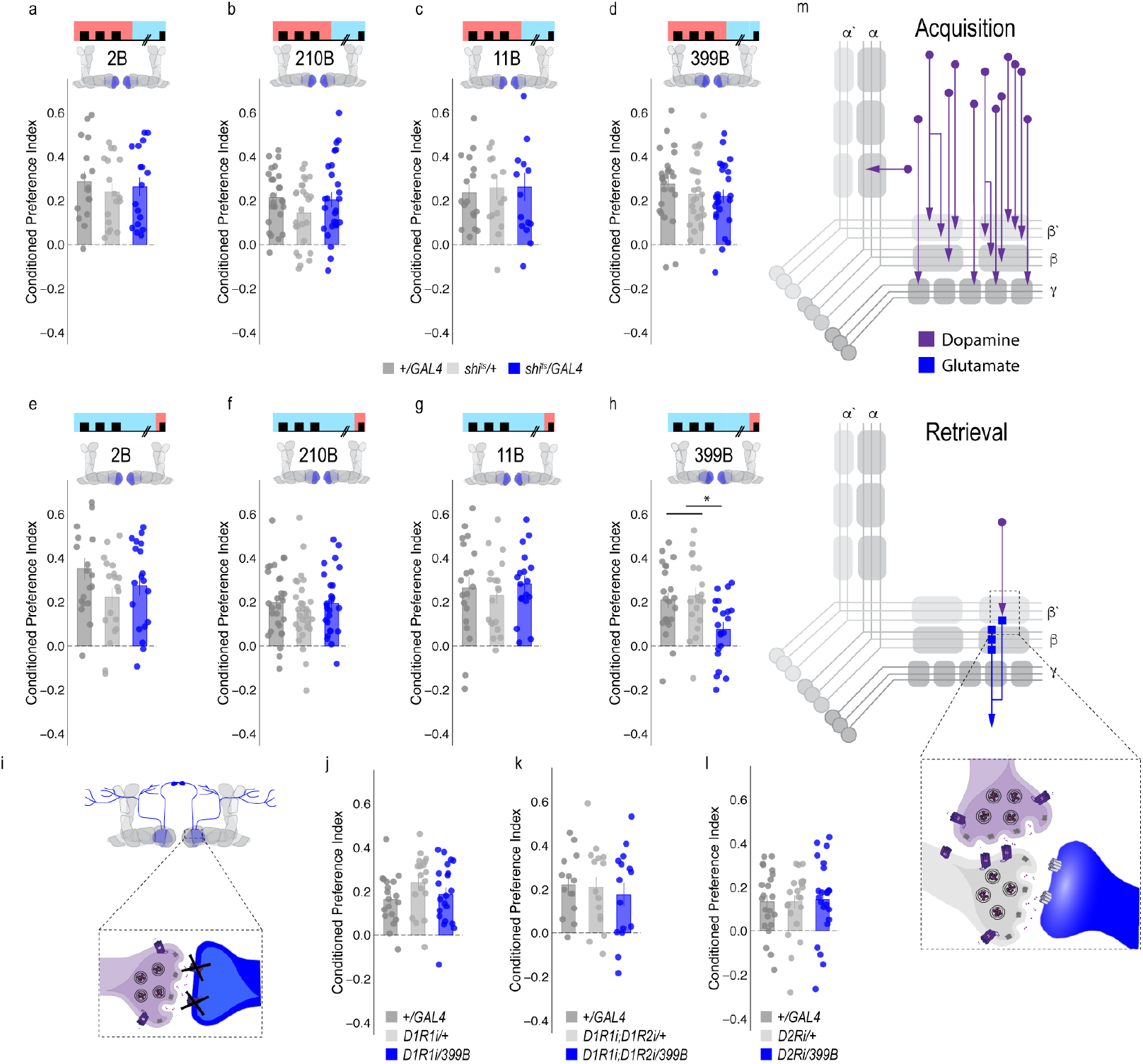
Memory expression during retrieval, but not acquisition, is dependent on a sparse population of glutamatergic MBONs. (a-d) Thermogenetic inactivation of glutamatergic MBONs innervating similar compartments to β’2a PAM dopaminergic neurons during acquisition did not disrupt encoding of alcohol associated preference. (e-h) However, inactivating neurotransmission during retrieval revealed the specific importance of MBON β2β’2a glutamatergic neuron for the expression of alcohol associated preference. (i-l) Knockdown of dopamine receptors in MBON β2β’2a did not disrupt alcohol associated preference suggesting that dopamine and MBON glutamate neurons interact indirectly via the MB. Bar graphs illustrate mean +/- standard error of the mean. Raw data are overlaid on bar graphs. Each dot is an n of 1, which equals approximately 60 flies (30 per odor pairing). One-way ANOVA with Tukey Posthoc was used to compare mean and variance. *p<0.01 (m) Updated model of circuits responsible for encoding alcohol associated preference during acquisition and circuits responsible for the expression of alcohol associated preference during retrieval. Retrieval circuits require specific subsets of dopaminergic neurons and a single MBON glutamatergic neuron innervating the β2’a compartment.

### Memory expression is dependent on a sparse subset of dopamine neurons

A hallmark of reward-encoding dopamine neurons is the gradual transfer of neural activation from reward delivery to cue onset during associative learning. During learning, dopaminergic activity shifts from responding to a reward during acquisition to the cue that predicts a reward during expression of the associated memory (Keiflin and Janak 2015, Schultz 2015, Schultz 2016). However, the circuit mechanisms underlying this shift and knowledge about whether all dopamine neurons do this, or whether a selective subset of dopamine neurons respond to the cue are unknown. It’s critical to understand this if we want to understand how drugs of abuse (alcohol) influence these circuits to result in such long-lasting, inflexible memories.

We therefore used a thermogenetic approach, as previously described, to temporarily inactivate neurotransmission in subsets of dopamine input neurons during retrieval, but not during acquisition or consolidation using a set of highly specific split-Gal4 lines. Strikingly, only inactivating dopaminergic neurons innervating β’2a compartment of the MB, using split-Gal4 line 109B, significantly reduced alcohol associated preference suggesting that these neurons are important for the expression of alcohol associated preference during retrieval (Figure 3). This suggests population encoding during acquisition shifts to sparse representation during memory expression.

### A dopamine-glutamate circuit regulates memory expression

Systems memory consolidation suggests that there are different circuits for memory acquisition and expression. Indeed, work in both fly and mammalian models suggest brain regions have a time limited role in systems consolidation. For example, recent work shows evidence of distinct acquisition and retrieval hippocampal circuits for contextually based fear memory (Roy et al 2017). Our next goal was to map the circuits through which the dopamine signal during retrieval drives the behavioral decision to move towards the paired cue.

To determine the circuit mechanisms that regulate a shift from dopaminergic population encoding during acquisition to sparse representation during retrieval, we tested the requirement of MB output neurons (MBONs) that aligned with the β’2a dopamine neurons required for memory expression. Inactivating the output of glutamatergic MBONs innervating similar compartments during acquisition, using 4 different split-Gal4 lines, did not significantly reduce alcohol associated preference (Figure 4A-D). However, similar inactivation during retrieval identified a single β2 β’2a glutamatergic MBON important for the expression of alcohol associated preference (Figure 4E-H).

Thus far, we have defined a putative microcircuit that consists of a single subset of 8-10 dopamine neurons innervating the β’2a MB compartment and a single glutamatergic MBON that also innervates the β’2a MB compartment (β2 β’2a) that are important for the expression of alcohol associated preference (Figure 4m). Previous work suggested that β’2a dopaminergic neurons were anatomically connected with β’2amp MBONs at the level of the MB, however, it was unclear which MBON β’2a dopaminergic neurons were synaptically connected (Lewis et al 2015). To confirm connectivity between β’2a dopaminergic neurons and β2 β’2a MBONs we used the recently developed anterograde transsynaptic labeling method trans-Tango to label the postsynaptic targets of the β’2a dopaminergic neurons (Talay, Richman et al. 2017 Figure 5a). Crossing split-Gal4 line *MB109B* with trans-Tango flies revealed α’β’ MB neurons as postsynaptic to β’2a dopaminergic neurons (Figure 5a). Interestingly, β’2mp MBON, and not β2 β’2a MBON were labeled as post synaptic to β’2a dopaminergic neurons suggesting that connections between β’2a dopaminergic neurons and β2 β’2a glutamatergic MBONs are not direct, but likely interact via intrinsic cholinergic MB neurons (Barnstedt, Owald et al. 2016).

**Fig. 5.**
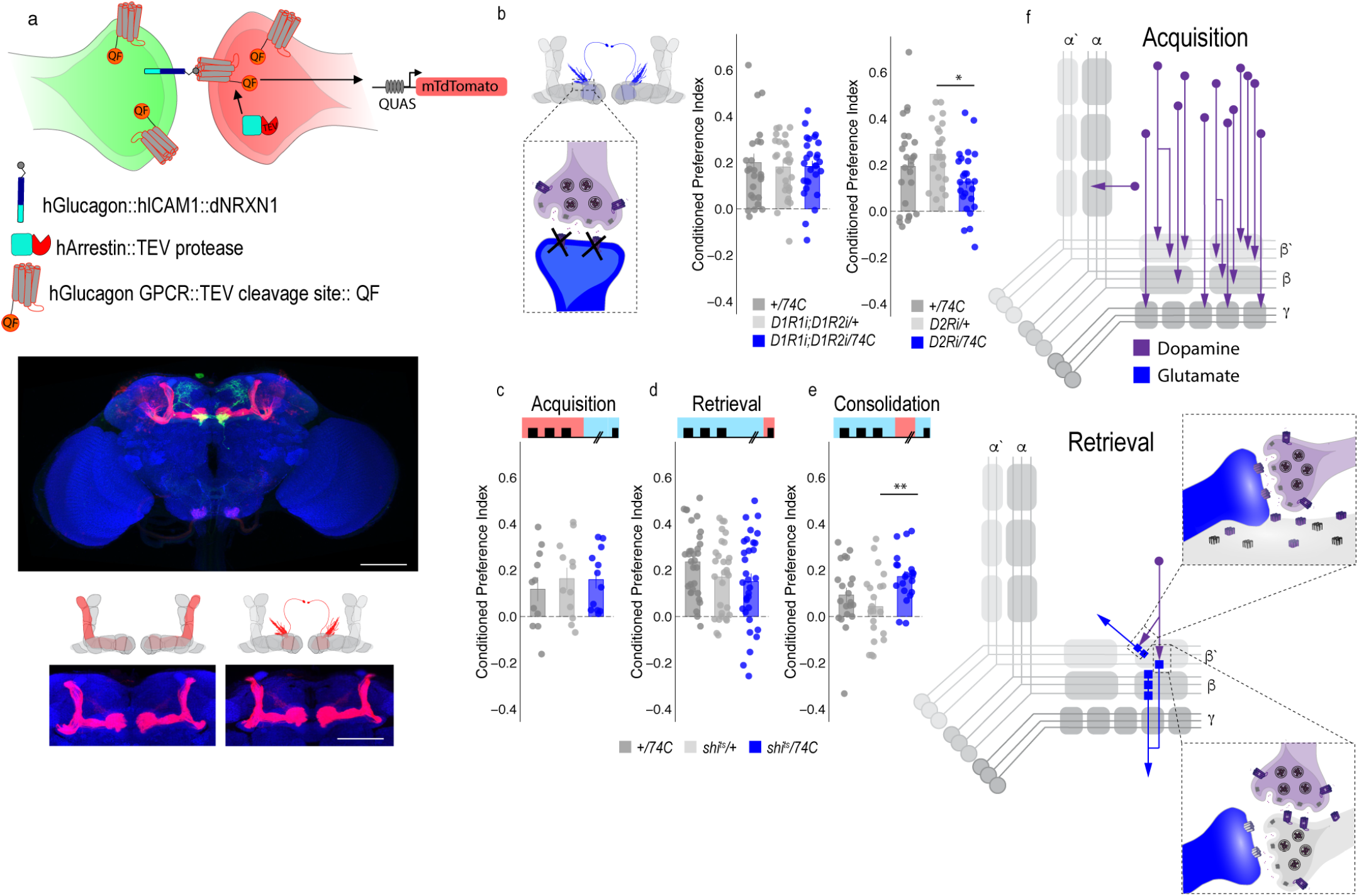
MBON β’2mp glutamatergic neuron is postsynaptic to β’2a PAM dopaminergic neuron and important for memory consolidation. (a) Schematic of *trans-*Tango (Talay, Richman et al. 2017). The GAL4/UAS system is used to express GFP and a transmembrane-tethered form of human glucagon localized to the presynaptic terminal via Neurexin1. All neurons have the potential to provide a read out of synaptic connectivity with the expression of a modified human glucagon receptor that has a TEV cleavage site and tethered QF. Postsynaptic neurons are genetically identified when glucagon activates its receptor, thereby recruiting Arrestin with a fused TEV protease to the receptor releasing QF to translocate into the nucleus and initiate mtdTomato expression from QUAS. Representative maximum projection confocal stacks reveal the α’/β’ MB lobe and MBON β’2mp neurons as postsynaptic to β’2a dopaminergic neurons. (b) Knockdown of D2Ri within MBON β2β’2a significantly reduced alcohol associated preference relative to UAS controls, but not GAL4 controls. (c-d) Thermogenetic inactivation of neurotransmission during acquisition or retrieval did not affect the expression of alcohol reward preference. (e)Thermogenetic inactivation during consolidation significantly increased alcohol reward preference relative to UAS controls, but not GAL4 controls. One-way ANOVA with Tukey Posthoc was used to compare mean and variance. Bar graphs illustrate mean +/- standard error of the mean. * p<0.05 **p<0.01 (f) Updated model of circuits responsible for encoding alcohol associated preference during acquisition and circuits responsible for the expression of alcohol associated preference during retrieval. Scale bar = 50 μm

To exclude the possibility of a false negative result, we confirmed the lack of functional connectivity between dopaminergic neurons and β2 β’2a using dopamine receptor RNAi (Supplementary Figure 5). RNAi against D1-like receptors or D2Rs in β2 β’2a neurons did not disrupt alcohol associated preference (Figure 4j-l). Combined, these data suggest the functional connectivity between β’2a dopaminergic neurons and the β2β’2a is not required for alcohol associated preference, but that expression of alcohol memories is occurring through a circuit consisting of MB cholinergic neurons, β’2a dopamine neurons and β2β’2a glutamate output neurons. Indeed, previous work from our lab reported the requirement of D2Rs in MB neurons for alcohol associated preference (Petruccelli, Feyder et al. 2018). A similar circuit motif was recently described in mice where dopamine regulated the release of acetylcholine from striatal tonically active interneurons on to glutamatergic neurons resulting in chronic decreases in corticostriatal activity during amphetamine withdrawal (Wang, Darvas et al. 2013). This suggests our findings describe a general circuit motif for exploring circuitry mechanisms of reward and motivation.

### A separate dopamine-glutamate circuit regulates memory consolidation

Our transsynaptic tracing method suggests a putative direct synaptic connection between β’2a dopamine neurons and β’2mp glutamatergic output neurons in regulating alcohol associated preference. We functionally validated the connection between β’2a dopamine neurons and β’2mp glutamatergic neurons using dopamine receptor RNAi lines (Figure 5b). Interestingly, decreasing levels of D2R, but not D1Rs, reduced alcohol associated preference (Fig 5b), suggesting a D2R-dependent pathway in MB that regulates alcohol memory.

Similar circuit motifs whereby dopamine directly synapses on glutamatergic neurons in the prefrontal cortex (PFC) have been reported in mammals and importantly, are altered by alcohol (Trantham-Davidson and Chandler 2015). These connections might support memory flexibility; however, it is unclear the mechanism by which this works. We hypothesized that neurons important for memory consolidation in our *Drosophila* MB circuit might modulate expression of memory, thereby influencing flexibility of memory. Previous work in *Drosophila* reported that activating β’2mp glutamatergic neurons promoted arousal (Sitaraman et al 2015). Thus, we reasoned that inactivating these neurons while flies normally sleep would further decrease arousal and facilitate memory consolidation. To confirm this hypothesis, we tested how activity of MB074C neurons affected alcohol memory as this line, which expresses highly in β’2mp, as well as β2 β’2a, and to a much less extent, γ5 β’2a (Aso, Hattori et al. 2014) aligns with the predicted β’2mp postsynaptic cells. Inactivating these subsets of glutamatergic neurons during acquisition or retrieval did not disrupt alcohol associated preference (Figure 5c, d). However, when these neurons were inactivated during the overnight consolidation, alcohol associated preference was enhanced relative to controls (Figure 5e). Together these data suggest that dopamine via β’2a neurons inhibit the β’2mp glutamatergic neuron via D2R receptors which leads to the expression of alcohol associated preference. In the absence of dopamine (Figure 3f) or D2R receptors (Figure 5b), preference is disrupted.

### Convergent microcircuits encode alcohol reward expression

The central role for the β’2mp in consolidation and memory malleability suggests that this region may integrate information from several circuits required for memory expression. Indeed, there is a wealth of examples in the literature of the systems balancing input from integrating neural circuits to drive goal directed behavior (Knudsen 2007, Buschman and Miller 2014, Hoke, Hebets et al. 2017). Previous anatomical studies predicted that β’2mp glutamatergic MBON and α’2 cholinergic MBON were synaptically connected (Aso, Hattori et al. 2014). trans-Tango confirmed that the β’2mp MBON is a postsynaptic target of the α’2 MBON (Figure 6a).

**Fig. 6.**
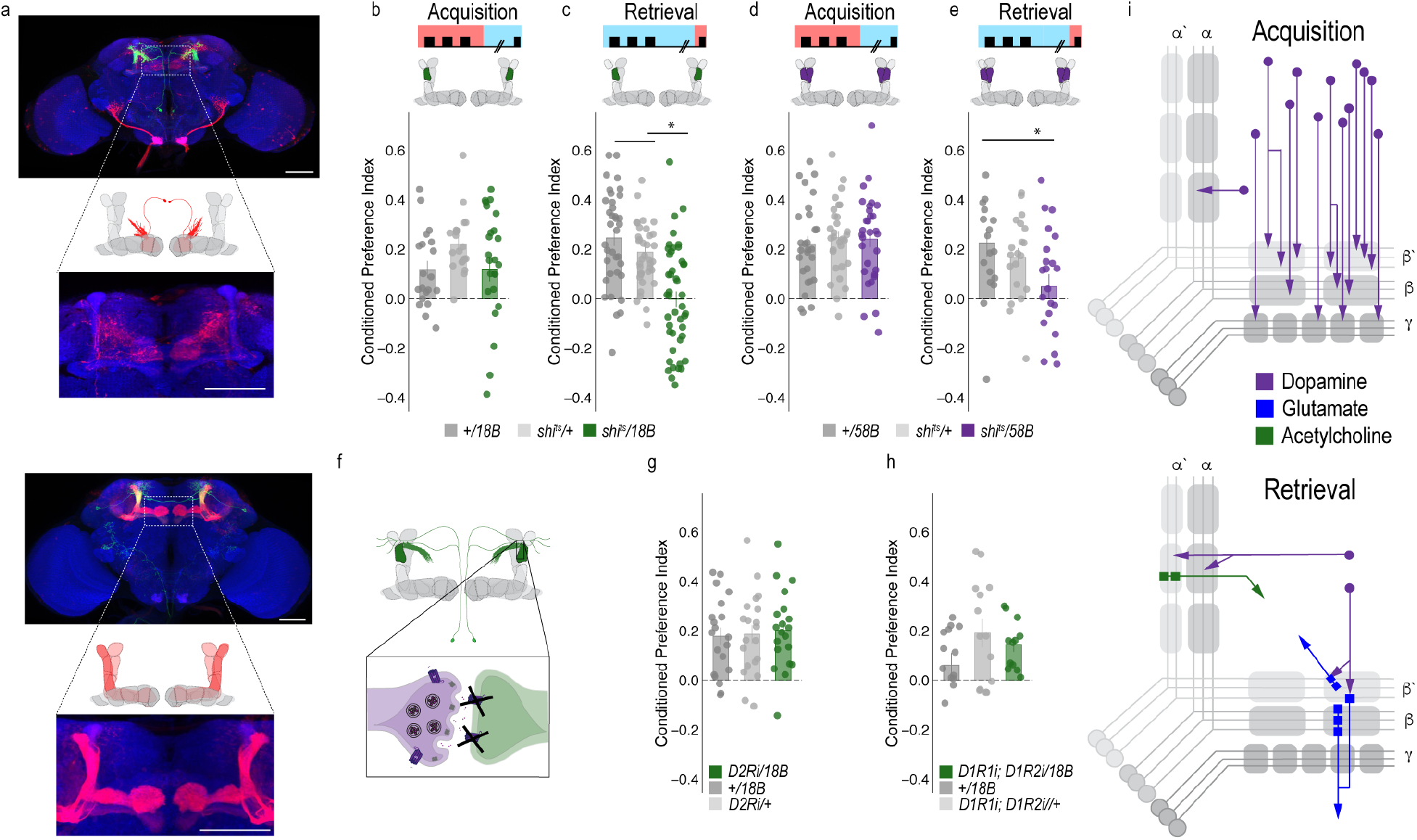
A microcircuit within the vertical lobe is important for alcohol associated preference. (a) *trans-*Tango experiments confirms that β’2mp glutamatergic MBON is postsynaptic to α’2 cholinergic neuron. (b-c) Thermogenetic inactivation of α’2 cholinergic neurons during acquisition did not affect the expression of alcohol associated preference, however, inactivation during retrieval significantly reduced preference. (d-e) Similarly, thermogenetic inactivation of α2α’2 dopaminergic neurons during acquisition did not affect the expression of alcohol associated preference, however, inactivation during retrieval significantly reduced preference. (f) *trans-*Tango experiments reveal that α2α’2 dopaminergic neurons are not synaptically connected to α’2 cholinergic MBON. (g-i) Knockdown of either D2R receptors or D1R receptors did not affect alcohol associated preference. (j) Updated model of circuits responsible for encoding alcohol associated preference during acquisition and circuits responsible for the expression of alcohol associated preference during retrieval. Scale bar = 50 μm

We previously showed that inactivating the α’2 cholinergic output throughout both memory acquisition and expression decreased alcohol associated preference (Aso, Sitaraman et al. 2014). To establish the specific temporal requirements of α ’2 MBON and determine whether its corresponding α 2 α ’2 dopaminergic input is necessary for alcohol associated preference, we thermogenetically inactivated neurotransmission during either acquisition or retrieval. Inactivating α’2 cholinergic MBONs or its corresponding α2 α’2 dopaminergic neurons during expression, but not acquisition of alcohol memory, significantly reduced alcohol associated preference (Figure 6b-e). The involvement of α2 α’2 dopaminergic neurons is particularly interesting because it part of a separate population of dopamine neurons, the paired posterior lateral 1 (PPL1) population, which is responsible for assigning negative valences to associated cues (Aso, Herb et al. 2012, Waddell 2013). This suggests that a second microcircuit in the vertical lobe, which converges onto the β’2mp neuron, is important for alcohol associated preference.

Interestingly, trans-Tango did not identify the α’2 cholinergic MBON as a postsynaptic target of α2 α’2 dopaminergic neuron. Similarly, RNAi against D1-like receptors or D2-like receptors did not disrupt alcohol associated preference (Figure 6f-h), suggesting that either these subsets of neurons are not directly connected or their direct connectivity is not required for alcohol associated preference. This type of connectivity is striking because it suggests that microcircuits important for the expression of memory converge on a neuron whose activity regulates consolidation, which might underlie memory flexibility. Future studies should address how the relative activity of β’2mp can augment memory expression.

### Alcohol memory expression circuits converge on a higher-order integration center

Thus far we have identified two MB microcircuits, one in the horizontal lobe that consists of PAM dopaminergic neurons and glutamatergic MBONs and one in the vertical lobe that consists of PPL1 dopaminergic neurons and cholinergic MBONs. Interestingly these two microcircuits converge on to a separate glutamatergic MBON (β’2mp) that is important for arousal and when inactivated, facilitates consolidation (Figure 6i). This suggests a circuit framework through which alcohol could shift memory from flexible to inflexible. However, it is still unclear how the MB and MBON activity drive goal directed behavior in the fly. Emerging models in the MB field suggest that MBON activity is pooled across compartments and that learning shifts the balance of activity to favor approach or avoidance (Owald and Waddell 2015). It remains unclear where this MBON activity converges.

In order to identify potential regions that integrated MBON activity, we used trans-Tango to map postsynaptic partners of α’2, β’2mp, and β2 β’2a MBONs. As mentioned previously, α’2 MBON labeled with MB018B-split-GAL4 identified β’2mp as a postsynaptic target. In a number of flies, γ5 β’2a was also identified as a target, however, this was less consistent. Interestingly, the dorsal regions of the FSB, specifically layers 4/5 or layer 6, were consistently identified as postsynaptic target of α’2 (Figure 7a, c). Strikingly both β’2mp and β2 β’2a also have synaptic connectivity with the dorsal regions of the FSB (Figure 7b, d). Together these data reveal the dorsal FSB as an intriguing convergent region downstream of the MB whose role in alcohol associated preference should be investigated further (Figure 7e).

**Fig. 7.**
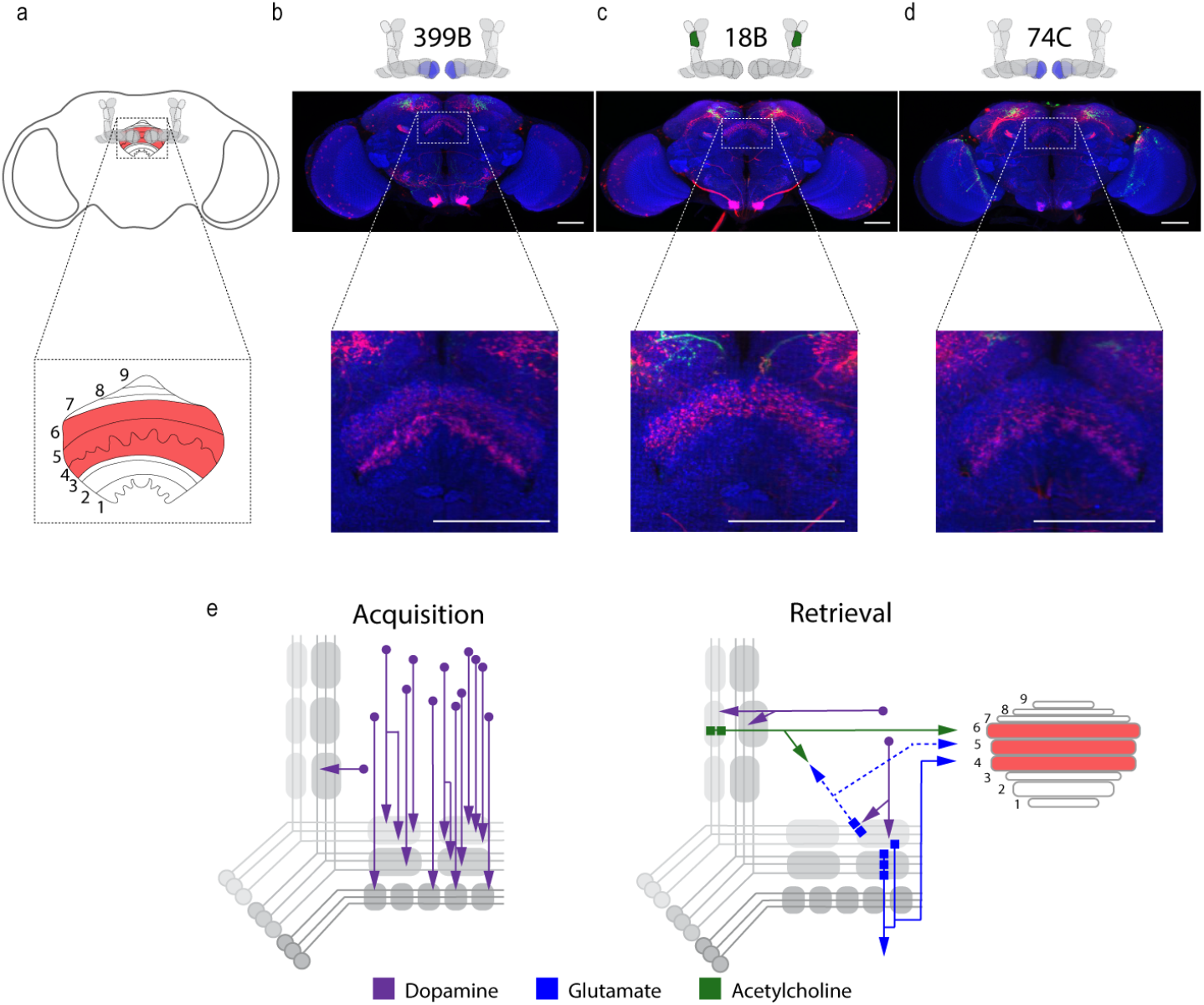
Circuits important for memory expression at retrieval converge on the dorsal FSB. (a) Schematic of fly brain with FSB and its layers highlighted. The FSB is a 9-layer structure (Wolff, Iyer et al. 2015), of which layers 4, 5, and 6 are targets. (b) Confocal stack of FSB highlighting the postsynaptic signal of β2β’2a MBON in the FSB. This MBON predominately targets layers 4, and 6. (c) Confocal stack of FSB highlighting the postsynaptic signal of α’2 MBON in the FSB. This MBON predominately targets layer 6. (d) Confocal stack of FSB highlighting the postsynaptic signal of β’2mp MBON in the FSB. This MBON predominately targets layer 4 and 5. (e) Updated model of circuits responsible for encoding alcohol associated preference during acquisition and circuits responsible for the expression of alcohol associated preference during retrieval. Model highlights the interconnectivity of the vertical and horizontal microcircuits and layers within the dorsal FSB where MBONs converge. Scale bar = 50 μm

## Discussion

A classic hallmark of addiction is the enduring propensity to relapse despite prolonged abstinence. Often relapse is driven by drug associated cues that evoke powerful and pervasive memories. Here we address the mechanisms underlying this persistence by investigating how these memories are encoded and expressed using a combination of behavioral, thermogenetic, in vivo calcium imaging and genetic transsynaptic tracing approaches in the fruit fly, *Drosophila melanogaster.*

Alcohol is a unique stimulus, because unlike natural rewards or punishers, it has both aversive and appetitive properties and yet the long-standing memories of intoxicating experiences are enduringly appetitive. Strikingly, the expression of this preference is not immediate. When flies are presented with a choice between the unpaired odor and the paired odor 30 minutes following acquisition of alcohol memory, flies avoid the paired odor and instead show a preference for cues not associated with alcohol (Kaun, Azanchi et al. 2011). Interestingly, cue associated avoidance switches to an enduring preference 24 hours later. The temporal nature of this valence switch suggests that, like mammals, the neural circuitry underlying the acquisition and expression of cue associated preference in flies is both anatomically complex, temporally distinct, and involves long-lasting changes to the circuitry mechanism.

Here we show that contrary to adaptive aversive or appetitive memories in flies (Liu, Placais et al. 2012, Masek, Worden et al. 2015, Yamagata, Ichinose et al. 2015, Yamagata, Hiroi et al. 2016), encoding alcohol associated preference is not dependent on a single subset of dopamine neurons or output neurons projecting from the MB, but a population of dopamine neurons whose activity emerges over the course of exposure to intoxicated doses of alcohol. We postulate that an increased requirement of dopamine neurons contributes to the stability of alcohol memory. Similarly, increasing the number of DA neurons that encode an aversive stimulus enhances how long memory lasts in *Drosophila* (Aso and Rubin 2016). This suggests a general rule where stability of memory is encoded by the number of dopamine neurons involved during acquisition. Most drugs of abuse initially increase dopamine levels beyond what is experienced during natural reward (Nutt, Lingford-Hughes et al. 2015, Volkow and Morales 2015, Kegeles, Horga et al. 2018). Our data suggests that this likely occurs, at least in part, via recruitment of additional dopamine neurons. Understanding the mechanism by which dopamine neurons are recruited may provide powerful insight into why memories for an intoxicating experience are so persistent.

In order to understand the mechanism of memory persistence, it is necessary to understand how dopamine encodes memories for cues associated with reward. In both primates and rodents, studies report a distinctive response profile whereby dopamine neuronal activity shifts its response from the presentation of a reward during learning to the presentation of a reward-predictive cue during expression (Keiflin and Janak 2015, Schultz 2016). However, given the heterogeneity of neurons in the VTA and the fact that these responses are typically observed using extracellular recordings, it is unclear how many and which dopamine neurons within the VTA and SNc exhibit this distinctive response to reward-predictive cues. Our intersectional genetic approach in *Drosophila* provided the temporal and spatial resolution to address this question. We found responses to reward-predictive cues were restricted to a small subset of dopamine neurons that were active during the initial alcohol presentation. These data suggest that cue responses are sparsely represented within a discrete subset of dopamine neurons found within the larger population of reward encoding dopamine neurons. Having identified a subset of alcohol associated cue-responsive dopamine neurons, we are now poised to further examine how the activity pattern of these neurons change with time and experience in vivo.

Additionally, we found cue-responsive dopamine neurons make direct connections with a glutamatergic neuron implicated in arousal (Sitaraman, Aso et al. 2015). Blocking this β’2mp neuron when flies normally sleep enhanced memory in a D2R-dependent manner. We propose that β’2a dopamine neurons inhibit β’2mp glutamate neuronal activity, thus permitting consolidation of alcohol associated preference. Strikingly, β’2a dopaminergic neurons were previously reported to inhibit the output of β’2amp MBONs to promote approach behaviors when flies were presented with conflicting aversive and appetitive odor cues (Lewis, Siju et al. 2015). Like other animals, flies find CO_2_ aversive, however, in the context of decaying fruit, flies often approach appetitive food odor cues despite the innately aversive CO_2_ odor cues. Lewis et al (2015) demonstrated β’2a dopaminergic neurons activity was required to overcome the innately aversive properties of CO_2_ and exhibit preference for these combined odor cues. The effects of β’2a dopamine neuronal inhibition, however, were not long lasting and thus responses to CO_2_ remained flexible outside the context of appetitive food odors. Here, the appetitive food odor, and consequently the activity of β’2a dopaminergic neurons, appears to act as an occasion setter, or a discriminatory stimulus that augments an animal’s response to a cue, thereby changing fly’s typical response to CO_2_ (Lewis, Siju et al. 2015). We speculate this neuron also resets the response to a cue associated with alcohol, which may be critical for overcoming the initial aversive properties of alcohol. We postulate that repeated intoxicating experiences change the dynamics of β’2a dopamine neurons during acquisition or consolidation in a way that create long term changes to the responsivity of the β’2mp MBON. This ultimately results in an inflexible memory circuit that supports alcohol associated preference.

Once acquired, distinct processes regulate consolidation and expression of enduring preferences for cues associated with alcohol intoxication. We found this process relies on a complex multilevel neural circuit that extends across brain regions to integrate information and consist of two converging microcircuits, each with its own subset of dopaminergic neurons and MBONs. These microcircuits emerge with time, are not necessary for the acquisition of these long-lasting preference associations, and converge on the β’2mp glutamate neuron (Figure 8). The β’2a dopamine neurons, which innervate the β’2mp glutamate neuron, make indirect connections through odor-coding MB Kenyon cells to the β2β’2a glutamate neurons. Similarly, the α2α’2 dopamine neurons make indirect connections with the α’2 cholinergic neuron, which also innervates the β’2mp neuron. The requirement for dopamine neurons implicated in encoding both reward and punishment in the expression of alcohol memory speaks to how seemingly conflicting events are integrated to produce an appropriate behavioral response.

**Fig. 8.**
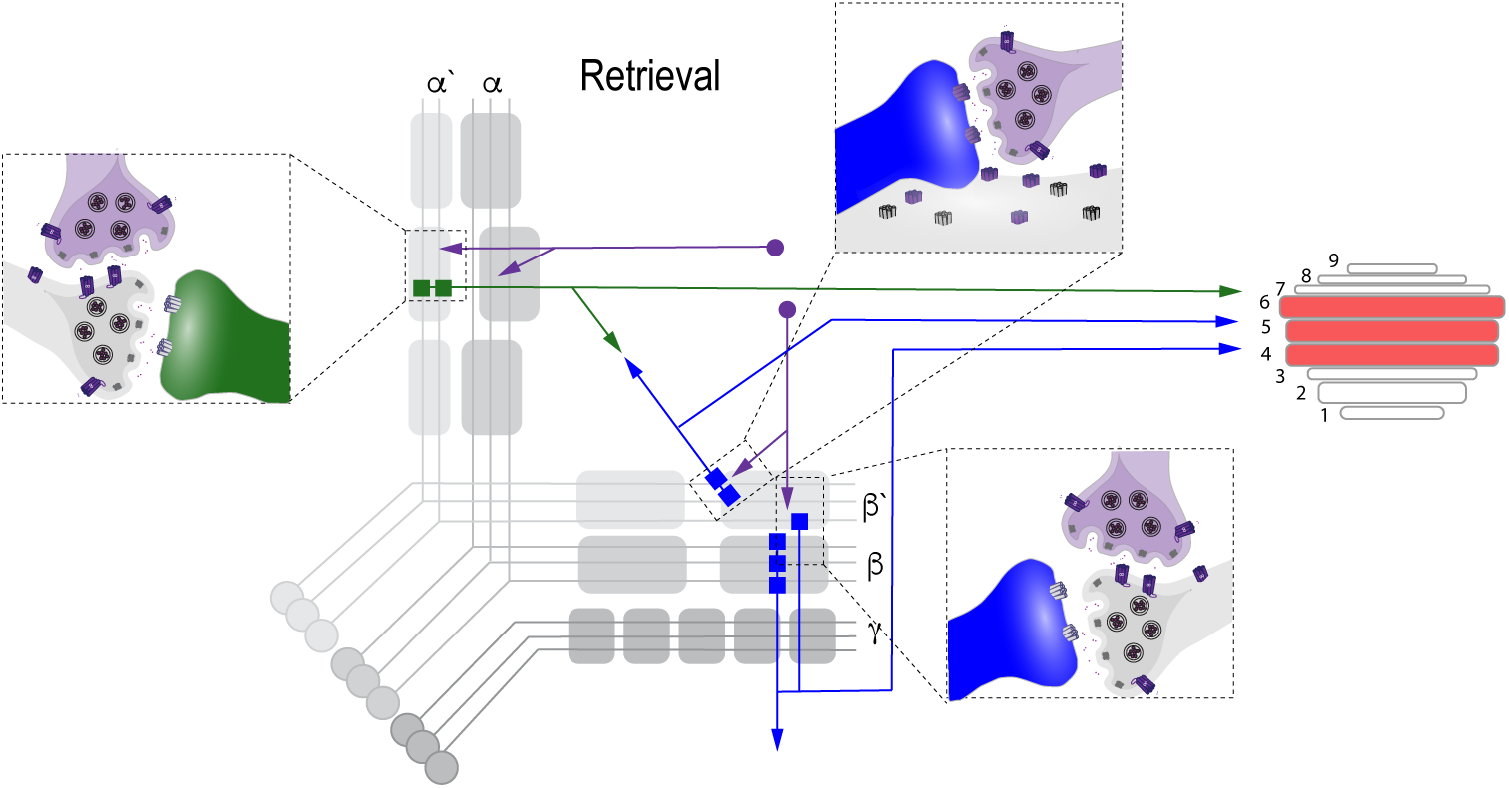
Proposed circuit important for the expression of alcohol associated preference at retrieval. The proposed circuit comprises two separate microcircuits, one in the vertical lobe and one in the horizontal lobe that converge on β’2mp MBON as well as layers with the dorsal FSB. The vertical lobe microcircuit includes a PPL1 α’2α2 dopaminergic neuron that have indirect connections with an α’2 cholinergic MBON via the MB. The horizontal lobe microcircuit includes a subset of PAM dopaminergic neurons (β’2a) that have direct connections with the β’2mp glutamatergic MBON and indirect connections with the β2β’2a glutamatergic MBON. Vertical and horizontal microcircuits converge on the β’2mp glutamatergic MBON which is important for arousal (Sitaraman, Aso et al. 2015) and layers 4, 5, and 6 of the FSB.

Our data suggest that activity of these β’2a and α’2 dopamine-MBON microcircuits is required for expression of alcohol associated preference. Unlike sucrose or shock memory formation where dopamine neurons inhibit the response of MBONs to elicit an appropriate behavioral response following training (Perisse, Yin et al. 2013, Aso, Sitaraman et al. 2014, Hige, Aso et al. 2015, Owald, Felsenberg et al. 2015, Aso and Rubin 2016, Cognigni, Felsenberg et al. 2018), our data suggest that activation of both the dopamine and output neurons in β’2a and β’2 compartment are required to initiate an alcohol seeking behavioral response. Because the β’2mp neuron is not required for expression of memory, it is likely that its output is integrated elsewhere in the brain to drive goal directed behaviors.

Further investigation of downstream targets of these microcircuits identified an additional convergent region: the dorsal layers of the FSB, specifically layers 4, 5, and 6. The identification of the FSB as a convergence region of MBONs is notable because a popular model proposes that learned behavioral responses in the fly are a consequence of pooled activity of mushroom body output neurons, which shifts the gain to either approach or avoidance behavior (Owald and Waddell 2015). Here we have identified one such structure that is an anatomical candidate for pooling MB output activity to drive learned behaviors. Interestingly, although the FSB has an established role in arousal and sleep, more recent work has defined its role in innate and learned nociceptive avoidance further supporting its role in integrating MB output activity (Hu, Peng et al. 2018). We hypothesize that output signals from the β2β’2a and α’2 neurons are integrated at the FSB to shift naïve response to cue-directed learned response. Compellingly, the β’2mp neuron also sends projections the FSB, presenting a circuit framework through which flexibility of memory could influence memory expression.

Together, these results provide valuable insight to the dynamic qualities of memory and how the long-lasting reward memories for alcohol intoxication are acquired, maintained, and expressed. We demonstrate that appetitive response for alcohol is encoded within a population of dopamine neurons, and expressed through microcircuits defined by subsets of these dopamine neurons. These microcircuits converge at several points; one through a neuron that regulates consolidation, and another at a downstream target implicated in integration of naïve and learned responses. This provides a new framework for how drugs of abuse regulate acquisition and expression of sensory memories, which ultimately results in a shift in behavioral response from malleable to inflexible.

## Materials and Methods

### Fly Strains

All *Drosophila melanogaster* lines were raised on standard cornmeal-agar media with tegosept anti-fungal agent and maintained at either 18°C or 21°C. For a list of fly lines used in the study, see Table 1. All *Drosophila melanogaster* lines used for trans-Tango were raised and maintained at 18°C in humidity-controlled chambers under 14/10 hr light/dark cycles on standard cornmeal-agar media with tegosept anti-fungal agent.

### Behavioral Experiments

Odor Preference Conditioning: For behavior experiments, male flies were collected 1-2 days post eclosion and were shifted from 21°C to 18°C, 65% humidity and placed on a 14/10 hr light/dark cycle. Odor conditioning was performed similar to Kaun et al. 2011. In short, groups of 30 males were trained in perforated 14ml culture vials filled with 1ml of 1% agar and covered with mesh lids. Training rooms were temperature and humidity controlled (65%). Training was performed in the dark with minimal red-light illumination and was preceded by a 20-minute habituation to the training chambers. Training chambers were constructed out of PlexiGlas (30 × 15 × 15 cm) (for details please refer to (Nuñez, Azanchi et al. 2018)). During habitation, humidified air (flow rate: 130) was streamed into the chambers. A single training session consisted of a 10 minute presentation of odor 1 (flow rate: 130), followed by a 10 minute presentation of odor 2 (flow rate 130) with 60% ethanol (flow rate 90: ethanol 60:air). Reciprocal training was performed simultaneously to ensure that inherent preference for either odor did not affect conditioning scores. For the majority of experiments odors used were 1:36 isoamyl alcohol and 1:36 isoamyl acetate in mineral oil, however, screen behavioral experiments used 1:36 isoamyl alcohol and 1:36 ethyl acetate in mineral oil. Vials of flies from Group 1 and Group 2 were age matched and paired according to placement in the training chamber. Pairs were tested simultaneously 24 hours later in the Y maze by streaming odor 1 and odor 2 (flow rate 10) in separate arms and allowing flies to walk up vials to choose between the two arms. A preference index was calculated by # flies in the Paired Odor Vial- # flies in the Unpaired Odor Vial)/ total # of flies that climbed. A conditioned preference index (CPI) was calculated by the averaging preference indexes from reciprocal groups. All data are reported as CPI. All plots were generated in RStudio.

Odor Sensitivity: Odor sensitivity was evaluated at restrictive temperatures (30°C). Odors used were 1:36 iso-amyl alcohol in mineral oil and 1:36 is-amyl acetate in mineral oil. Groups of 30 naïve males were presented with either an odor (flow rate 10) or air streamed through mineral oil in opposite arms of the Y. Preference index was calculated by # flies in odor Vial- # flies in air vial)/ total # flies that climbed for each individual odor.

Ethanol Sensitivity: Ethanol sensitivity was evaluated in the recently developed flyGrAM assay (Scaplen and Mei et al 2019). Briefly, for thermogenetic inactivation, 10 flies were placed into arena chambers and placed in a 30°C incubator for 20 minutes prior to testing. The arena was then transferred to a preheated (30°C) light sealed box and connected to a vaporized ethanol/humidified air delivery system. Flies were given an additional 15 minutes to acclimate to the box before recordings began. Group activity was recorded (33 frames/sec) for five minutes of baseline, followed by 10 minutes of ethanol administration and five minutes of following ethanol exposure. Activity was binned by 10 seconds and averaged within each genotype. Mean group activity is plotted as a line across time with standard error of the mean overlaid. All activity plots were generated in RStudio.

### trans-Tango Immunohistochemistry and Microscopy

Experiments were performed according to the published FlyLight protocols with minor modifications. Briefly, either adult flies that are 15-20 days old were cold anaesthetized on ice, de-waxed in 70% ethanol dissected in cold Schneider’s Insect Medium (S2). Within 20 minutes of dissection, tissue was incubated in 2% paraformaldehyde (PFA) in S2 at room temperature for 55 minutes. After fixation, samples were rinsed with phosphate buffered saline with 0.5% Triton X-100 (PBT) and washed 4 times for 15 minutes at room temperature. Following PBT washes, PBT was removed and samples were incubated in SNAP substrate diluted in PBT (SNAP-Surface649, NEB S9159S; 1:1000) for 1 hour at room temperature. Samples were then rinsed and washed 3 times for 10 minutes at room temperature and then blocked in 5% heat-inactivated goat serum in PBT for 90 minutes at room temperature and incubated with primary antibodies (Rabbit α-GFP Polyclonal (1:1000), Life Tech #A11122, Rat α-HA Monoclonal (1:100), Roche #11867423001) for two overnights at 4°C. Subsequently, samples were rinsed and washed 4 times for 15 minutes in 0.5% PBT and incubated in secondary antibodies (Goat α-Rabbit AF488 (1:400), Life Tech #A11034, Goat α-Rat AF568 (1:400), Life Tech #A11077) diluted in 5% goat serum in PBT for 2-3 overnights at 4°C. Samples were then rinsed and washed 4 times for 15 minutes in 0.5% PBT at room temperature and prepared for DPX mounting. Briefly, samples were fixed a second time in 4% PFA in PBS for 4 hours at room temperature and then washed 4 times in PBT at room temperature. Samples were rinsed for 10 minutes in PBS, placed on PLL-dipped cover glass, and dehydrated in successive baths of ethanol for 10 minutes each. Samples were then soaked 3 times in xylene for 5 minutes each and mounted using DPX. Confocal images were obtained using a Zeiss, LSM800 with ZEN software (Zeiss, version 2.1) with auto Z brightness correction to generate a homogeneous signal where it seemed necessary, and were formatted using Fiji software (http://fiji.sc).

### Dopamine Immunohistochemistry and Microscopy

Groups of flies were exposed to either 10 minutes of air or 10 minutes of ethanol and dissected within 15 minutes of exposure on ice. Immunohistochemistry was performed according to (Cichewicz, Garren et al. 2017). With 15 minutes of dissection, tissue was transferred to fix (1.25% glutaraldehyde in 1% PM) for 3-4 hours at 4°C. Tissue was subsequently washed 3 times for 20 minutes in PM and reduced in 1% sodium borohydride. Then the tissue was washed 2 times for 20 minutes before a final wash in PMBT. Tissue was blocked in 1% goat serum in PMBT overnight at 4°C and incubated in primary antibody (Mouse anti-dopamine (1:40) Millipore Inc, #MAB5300) for 48 hours at 4°C. Following primary antibody incubation, tissue was washed 3 times in PBT for 20 minutes at room temperature and incubated in secondary antibody (Goat anti mouse 488 (1:200 in PBT) Thermo #A11029) for 24 hours at 4°C. The following day tissue was washed 2 times for 20 minutes in PBT and then overnight in fresh PBT. Tissue was rinsed quickly in PBS, cleared in FocusClear and mounted in MountClear (Cell Explorer Labs). Confocal images were obtained using a Zeiss, LSM800 with ZEN software (Zeiss, version 2.1). Microscope settings were established using ethanol tissue before imaging air and ethanol samples.

### Dopamine fluorescence analysis

Fluorescence was quantified in Fiji (Schindelin, Arganda-Carreras et al. 2012) using Segmentation Editor and 3D Manager (Ollion, Cochennec et al. 2013). In segmentation editor ROIs were defined using the selection tool brush to outline the MB in each slice and also outside a background region immediately ventral to MB that lacked defined fluorescent processes. 3D ROIs of the MB and control region were created by interpolating across slices. Geometric and intensity measurements were calculated for each ROI in 3D Manager and exported to CSV files. Integrated density for each ROI was normalized by the integrated density of control regions. Average integrated density for air and ethanol exposures are reported. All fluorescence quantifications were performed by a blinded experimenter.

### Calcium Imaging Protocol and Analysis

To express GCaMP6m in PAM neurons, UAS-GCaMP6m virgin female flies were crossed to male flies containing the R58E02-GAL4 driver. As previously mentioned, all flies were raised on standard cornmeal agar food media with tegosept anti-fungal agent and maintained on a 14/10-hour light/dark cycle at 24°C and 65% humidity.

#### Fly Preparation

Male flies were selected for imaging six days post-eclosion. Flies were briefly anesthetized on ice to transfer and fix to an experimental holder made out of heavy-duty aluminum foil. The fly was placed into an H-shaped hold cut out of the foil and glued in place using epoxy (5-min Epoxy, Devcon). The head was tilted about 70° to remove the cuticle from the back of the fly head. All legs were free to move, the proboscis and antenna remained intact and unglued. Once the epoxy was dry, the holder was filled with *Drosophila* Adult Hemolymph-Like Saline (AHLS). The cuticle was removed using a tungsten wire (Roboz Surgical Instruments Tungsten Dissecting Needle, .125 mm, Ultra Fine (Pk 10)) and forceps #5. The prepared fly in its holder was positioned on a customized stand underneath the two-photon scope. The position of the ball and the stream delivery tubes were manually adjusted to the fly’s position in the holder.

#### Imaging paradigm

Calcium imaging recordings were performed with a two-photon resonance microscope (Scientifica). Fluorescence was recorded from the PAM neurons innervating the mushroom body for a total duration of 80 to 95 minutes. The first 10 minutes the fly was presented an air stream, followed by 10 minutes of isoamyl alcohol. The fly was then presented with 10-minutes of isoamyl alcohol paired with ethanol followed by 50 minutes of streaming air. To avoid bleaching effects and to match the higher resolution imaging properties, the recording was not throughout the entire paradigm but spaced with imaging intervals of 61.4 seconds. Recordings were performed using SciScan Resonance Software (Scientifica). The laser was operated at 930 nm wavelength at an intensity of 7.5-8mW. Images were acquired at 512 × 512-pixel resolution with an average of 30.9 frames per second. Recordings lasted 1900 frames which equals 61.5 seconds. Recordings were performed at 18.5°C room temperature and 59% humidity.

#### Imaging analysis

Data were registered, processed, and extracted using a custom matlab GUI (Chris Deister github.com/cdesiter). Calcium image files (.tiff) comprising of 1900 frames taken at 30.94 frames per second rate (61.4 seconds), were initially averaged every 5 frames to downsize the .tiff image files to 380 frames. Image files were then aligned and registered in X-Y using a 15-50 frame average as a template. ROIs were constructed over the MB lobes using non-negative matrix factorization to identify active regions and then subsequently segmented to create the ROIs. Fluorescence values were extracted from identified ROIs and ΔF/F_o_ measurements were created using a moving-average of 75 frames to calculate the baseline fluorescence (F_o_). Average fluorescence traces across flies (n = 6) were visualized using ggplot in R studio. Fiji (Schindelin, Arganda-Carreras et al. 2012) was used to construct heat-maps visualizing calcium activity. Calcium image files were summated across 1900 frames to create Z-projections. A heat gradient was used to visualize calcium activity magnitude.

### qRT-PCR

qRT-PCR methods have been described previously (Petruccelli, Feyder et al. 2018). In brief, total RNA was extracted from approximately 100 heads using Trizol (Ambion, Life Technologies) and treated with DNase (Ambion DNA-Free Kit). Equal amounts of RNA (1 μg) were reverse-transcribed into cDNA (Applied Biosystems) for each of the samples. Then, Biological (R3) and technical (R2) replicates were analyzed with Sybr Green Real-Time PCR (BioRad, ABI PRISM 7700 Sequence Detection System) performed using the following PCR conditions: 15 s 95°C, 1 min 55°C, 40x. Primer sequences can be found in Supplementary Table 4. Across all samples and targets, Ct threshold and amplification start/stop was set to 0.6 and manually adjusted, respectively. All target genes were initially normalized to CG13646 expression for comparative ΔCt method analysis, then compared to control genotype to assess fold enrichment (ΔΔ Ct method).

## Declaration of Interests

The authors declare no conflict of interest.

This article contains supporting information online.

## Acknowledgments

This work was supported by the Smith Family Award Program for Excellence in Biomedical Research, United States, the Israel Binational Science Foundation Start Up Grant 2015005, the Carney Institute for Brain Science Center for Biomedical Research Excellence “Center for Nervous System Function” (NIGMS P20GM103645), NIAAA (R01AA024434), and the Rhode Island Foundation Medical Research Fund 20144133. We thank all of the Kaun lab members for fruitful discussions and feedback on earlier versions of this manuscript. We also thank the *Drosophila* community, particularly Yoshinori Aso and Gerry Rubin, the Bloomington Stock Center, and the Vienna *Drosophila* RNAi Center for sharing fly stocks. Yoshinori Aso also provided confocal stacks of MB042B and MB040B used for cell counting. We thank Thomas Boudier and Stefan Helfrich for helpful software advice, Jay Hirsh for reagents and advice, and Christopher Diester and other members of the Moore lab for calcium analysis guidance. Finally, we thank John McGeary, Tara White, and Daniel Dombeck for helpful comments on earlier version of this manuscript. BioRxiv Template from the Finkelstein Lab was used for formatting this manuscript (https://github.com/finkelsteinlab/BioRxiv-Template).

