## Supplemental Figures and Tables for "Circuits that encode and predict alcohol associated preference"

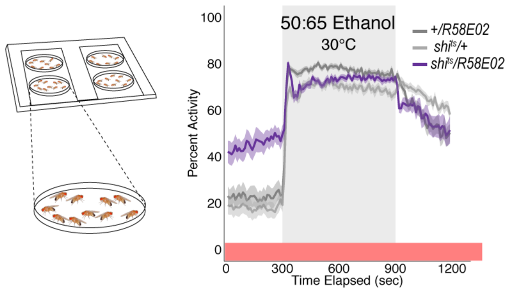


Supplementary Figure 1. Inactivation of PAM neurons do not affect alcohol induced activity suggesting that a decrease in preference is encoded independently from the amount of activity animals exhibit while intoxicated (Figure 1E). (a) Schematic of flyGrAM. Groups of ten male flies were placed into four behavioral chambers. Flies were exposed to five minutes of air, following by ten minutes of ethanol, and lastly 5 minutes of air. Group activity of flies was recorded at 33 frames per second.

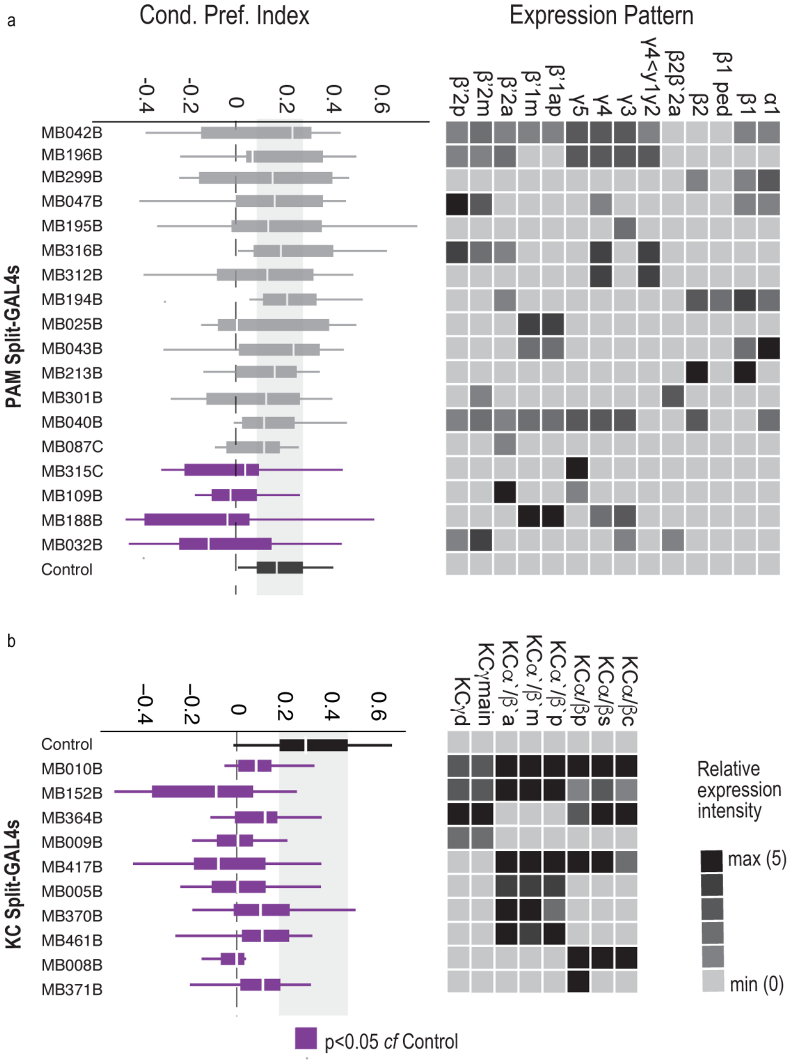


Supplementary Figure 2. (a) Subsets of dopamine neurons were inactivated during acquisition and retrieval using 18 specific PAM Split-GAL4 lines. Four lines, 315C, 109B, 188B, and 32B, which predominately express in the medial aspect of the MB resulted in significant decreases in preference for alcohol associated cues. (b) Subsets of Kenyon cells were inactivated during acquisition and retrieval using 10 specific Kenyon cell Split-GAL4 lines. All 10 lines resulted in significant decreases in preference for alcohol associated cues.

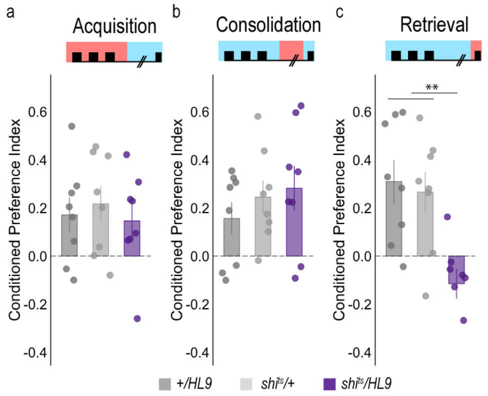


Supplementary Figure 3. Subsets of PAM neurons are required for retrieval, but not acquisition or consolidation. (a) Inactivating subsets of PAM neurons during acquisition using the HL9-GAL4 driver, which is not a split-GAL4 line, did not disrupt alcohol associated preference (Claridge-Chang, Roorda et al. 2009). (b) Inactivating subsets of PAM neurons during consolidation, defined as the overnight period between acquisition and retrieval did not disrupt alcohol associated preference. (c) However, inactivating the same subsets of PAM neurons during retrieval significantly disrupted alcohol associated preference. ** p<0.01

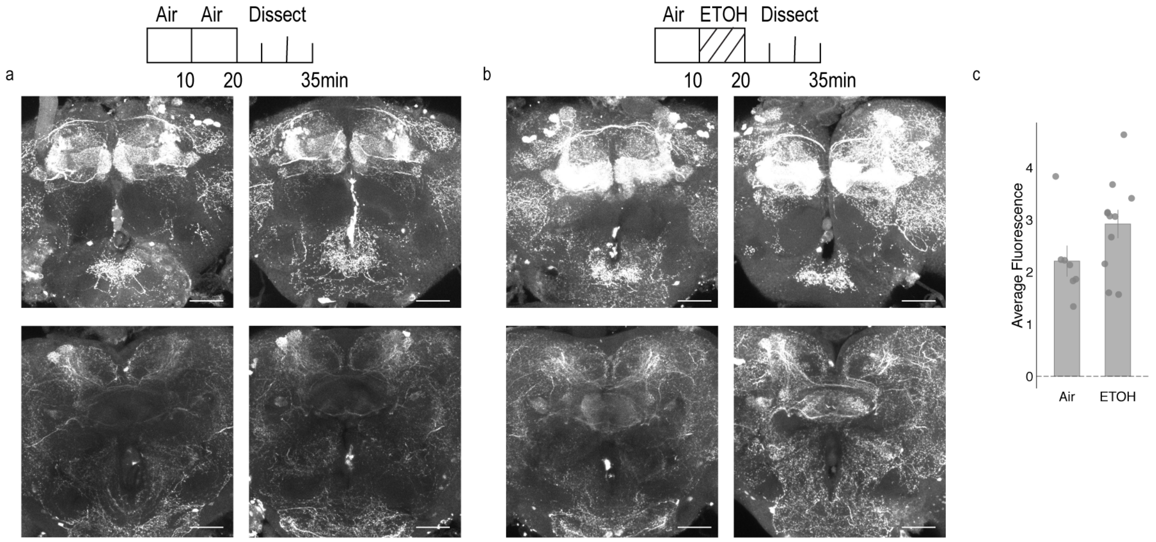


Supplementary Figure 4. Dopamine staining within the brain following 10 minutes of air or 10 minutes of ethanol. Left panel illustrates air condition. Flies were habituated for 10 minutes in the behavior box followed by an additional 10 minutes of air. Right panel illustrates ethanol condition. Flies were habituated for 10 minutes in the behavior box followed by 10 minutes of ethanol. In both conditions, the top panel illustrates staining with the MB. Maximum intensity z stacks were collected from the start of the gamma lobe to the end of the $\alpha/\beta$, $\alpha`\beta`$ lobes. Each stack consists of approximately 20 µm slices. Bottom panel illustrates staining within the central complex, predominately the FSB. Maximum intensity z stacks were collected from the start of the EB to the end of the FSB. Each stack consists of approximately 20 1um slices. Bar graphs illustrate mean +/- standard error of the mean. Raw data are overlaid on bar graphs. Each dot is 1 fly. One-way ANOVA was used to compare mean and variance.

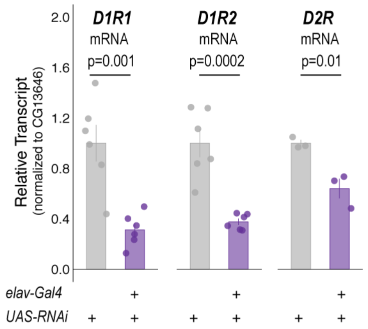


Supplementary Figure 5. Quantitative RT-PCR on whole heads of flies expressing respective *RNAi* with pan-neuronal *elav-Gal4*. Bar graphs illustrate mean +/- standard error of the mean. Raw data are overlaid on bar graphs. One-way ANOVA was used to compare mean and variance with Tukey Posthoc compared to experimental.

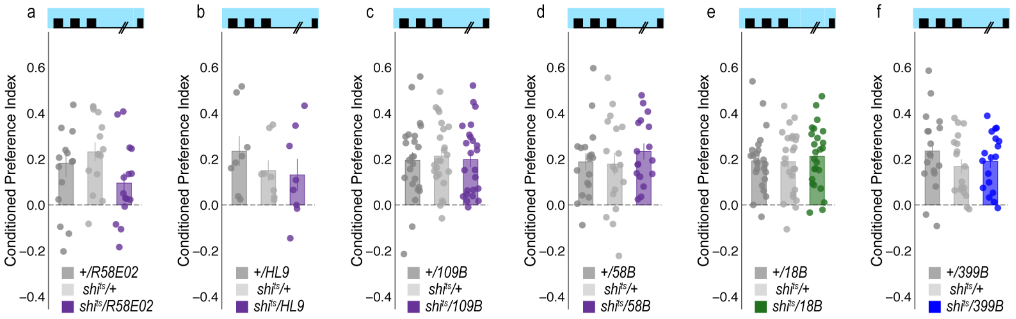


Supplementary Figure 6. Temperature controls for genotypes that showed decreases in alcohol associated preference. Flies were trained and tested at permissive temperatures (20°C) and exhibited normal alcohol associated preference for pair odor cues under these conditions.

| UAS-mcd GFP | | | |
| --- | --- | --- | --- |
| **40B** | **42B** | **HL9** | **R5802** |
| n=6 | n=6 | n=6 | n=4 |
| 44.83 ±5.26 | 41.00 ±1.37 | 60.10 ±07.54* | 101 ±3.43 |

Supplementary Table 1. PAM dopamine cell counts per hemisphere. *HL9 numbers from Claridge-Chang et al. 2009 (Claridge-Chang, Roorda et al. 2009)

| | **Odor1** | | | **Odor 2** | | |
| --- | --- | --- | --- | --- | --- | --- |
| **GAL4 lines** | **+/GAL4** | **+/UAS** | **GAL4/UAS** | **+/GAL4** | **+/UAS** | **GAL4/UAS** |
| R58E02 | 0.73 ±0.05 | 0.83 ±0.06 | 0.92 ±0.04 | 0.36 ±0.05 | 0.51 ±0.11 | 0.66 ±0.07 |
| HL9 | 0.53 ±0.11 | 0.49 ±0.08 | 0.62 ±0.08 | 0.49 ±0.10 | 0.44 ±0.13 | 0.33 ±0.07 |
| MB109B | 0.71 ±0.06 | 0.68 ±0.05 | 0.80 ±0.04 | 0.37 ±0.05 | 0.67 ±0.03 | 0.68 ±0.06 |
| MB058B | 0.69 ±0.04 | 0.78 ±0.04 | 0.90 ±0.02 | 0.56 ±0.04 | 0.57 ±0.04 | 0.67 ±0.03 |
| MB399B | 0.67 ±0.06 | 0.62 ±0.05 | 0.92 ±0.02 | 0.49 ±0.05 | 0.59 ±0.03 | 0.67 ±0.04 |
| MB018B | 0.65 ±0.06 | 0.74 ±0.06 | 0.57 ±0.04 | 0.47 ±0.07 | 0.41 ±0.09 | 0.48 ±0.04 |

Supplementary Table 2. Odor Controls at 30C. Odor 1: Isoamyl Acetate or Ethyl Acetate. Odor 2: Isoamyl Alcohol. Naïve flies were presented with either odor 1 vs air or odor 2 vs air in the Y maze.

| Figure | Experiment | n | Statistical Test | Result | Post-hoc | Result |
|---|---|---|---|---|---|---|
| 1c | Acquisition | +/R58E02 (n=23)<br>+/shit$^{s}$ (n=23)<br>shi$^{ts}$/R58E02 (n=23) | One-way ANOVA | F(2, 66)=5.355, p=0.007 | Tukey | +/ shi$^{ts}$ vs +/R58E02 p=0.87<br>+/ shi$^{ts}$ vs shi$^{ts}$/R58E02 p=0.009<br>+/R58E02 vs shi$^{ts}$/R58E02 p=0.04 |
| 1c | Consolidation | +/R58E02 (n=13)<br>+/shit$^{s}$ (n=15)<br>shi$^{ts}$/R58E02 (n=13) | One-way ANOVA | F(2,38)=5.964 | Tukey | +/ shi$^{ts}$ vs +/R58E02 p=0.004<br>+/ shi$^{ts}$ vs shi$^{ts}$/R58E02 p=0.18<br>+/R58E02 vs shi$^{ts}$/R58E02 p=0.26 |
| 1c | Retrieval | +/R58E02 (n=25)<br>+/shit$^{s}$ (n=24)<br>shi$^{ts}$/R58E02 (n=25) | One-way ANOVA | F(2,71)=5.707, p=0.005 | Tukey | +/ shi$^{ts}$ vs +/R58E02 p=0.65<br>+/ shi$^{ts}$ vs shi$^{ts}$/R58E02 p=0.05<br>+/R58E02 vs shi$^{ts}$/R58E02 p=0.005 |
| 1f | Calcium Imaging | GCaMP6m/R58E02 (n=6) | Repeated Measures ANOVA | F(3,15)=5.380, p=0.002 | Bonferroni | Odor Epoch 1 vs Odor + Ethanol Epoch 1 p=0.032<br>Odor Epoch 2 vs Odor + Ethanol Epoch 2 p=0.224<br>Odor Epoch 3 vs Odor + Ethanol Epoch 3 p=0.121<br>Odor Epoch 4 vs Odor + Ethanol p=0.004 |
| 2a | Acquisition | +/40B (n=23)<br>+/shit$^{s}$ (n=23)<br>shi$^{ts}$/40B (n=21) | One-way ANOVA | F(2,64)=1.262, p=0.39 | N/A | N/A |
| 2b | Acquisition | +/42B (n=24)<br>+/shit$^{s}$ (n=24)<br>shi$^{ts}$/42B (n=23) | One-way ANOVA | F(2,68)=0.995, p=0.38 | N/A | N/A |
| 2c | Acquisition | +/188B (n=10)<br>+/shit$^{s}$ (n=11)<br>shi$^{ts}$/188B (n=12) | One-way ANOVA | F(2,30)=0.084, p=0.92 | N/A | N/A |
| 2d | Acquisition | +/32B (n=22)<br>+/shit$^{s}$ (n=21)<br>shi$^{ts}$/32B (n=20) | One-way ANOVA | F(2,60)=1.52, p=0.23 | N/A | N/A |
| 2e | Acquisition | +/301B (n=27)<br>+/shit$^{s}$ (n=27)<br>shi$^{ts}$/301B (n=27) | One-way ANOVA | F(2,78)=0.389, p=0.68 | N/A | N/A |
| 2f | Acquisition | +/109B (n=24)<br>+/shit$^{s}$ (n=24)<br>shi$^{ts}$/109B (n=24) | One-way ANOVA | F(2,69)=0.091, p=0.91 | N/A | N/A |
| 2g | Acquisition | +/87C (n=23)<br>+/shit$^{s}$ (n=23)<br>shi$^{ts}$/87C (n=23) | One-way ANOVA | F(2,51)=0.663, p=0.52 | N/A | N/A |
| 2h | Acquisition | +/315C (n=20)<br>+/shit$^{s}$ (n=23)<br>shi$^{ts}$/315C (n=24) | One-way ANOVA | F(2,64)=0.24, p=0.79 | N/A | N/A |
| 3a | Retrieval | +/40B (n=7)<br>+/shit$^{s}$ (n=6)<br>shi$^{ts}$/40B (n=7) | One-way ANOVA | F(2,17)=2.43, p=0.12 | N/A | N/A |
| 3b | Retrieval | +/42B (n=16)<br>+/shit$^{s}$ (n=16)<br>shi$^{ts}$/42B (n=14) | One-way ANOVA | F(2,68)=0.995, p=0.38 | N/A | N/A |

| | | | | | | |
|---|---|---|---|---|---|---|
| 3c | Retrieval | +/188B (n=24)<br>+/shit$^{s}$ (n=27)<br>shi$^{ts}$/188B (n=25) | One-way ANOVA | F(2,73)=0.044, p=0.96 | N/A | N/A |
| 3d | Retrieval | +/32B (n=6)<br>+/shit$^{s}$ (n=6)<br>shi$^{ts}$/32B (n=6) | One-way ANOVA | F(2,15)=1.226, p=0.32 | N/A | N/A |
| 3e | Retrieval | +/301B (n=23)<br>+/shit$^{s}$ (n=24)<br>shi$^{ts}$/301B (n=24) | One-way ANOVA | F(2,78)=0.389, p=0.68 | N/A | N/A |
| 3f | Retrieval | +/109B (n=20)<br>+/shit$^{s}$ (n=24)<br>shi$^{ts}$/109B (n=24) | One-way ANOVA | F(2,65)=14.18, p= 7.78x10^-6 | Tukey | +/ shi$^{ts}$ vs +/109B p=0.07<br>+/ shi$^{ts}$ vs shi$^{ts}$/109B p=0.007<br>+/109B vs shi$^{ts}$/109B p=0.000005 |
| 3g | Retrieval | +/87C (n=20)<br>+/shit$^{s}$ (n=21)<br>shi$^{ts}$/87C (n=22) | One-way ANOVA | F(2,60)=0.266, p=0.77 | N/A | N/A |
| 3h | Retrieval | +/315C (n=20)<br>+/shit$^{s}$ (n=19)<br>shi$^{ts}$/315C (n=20) | One-way ANOVA | F(2,56)=0.109, p=0.90 | N/A | N/A |
| 4a | Acquisition | +/2B (n=16)<br>+/shit$^{s}$ (n=17)<br>shi$^{ts}$/2B (n=17) | One-way ANOVA | F(2,47)=0.31, p=0.73 | N/A | N/A |
| 4b | Acquisition | +/210B (n=26)<br>+/shit$^{s}$ (n=26)<br>shi$^{ts}$/210B (n=27) | One-way ANOVA | F(2,76)=1.59, p=0.21 | N/A | N/A |
| 4c | Acquisition | +/11B (n=17)<br>+/shit$^{s}$ (n=15)<br>shi$^{ts}$/11B (n=15) | One-way ANOVA | F(2,44)=0.09, p=0.92 | N/A | N/A |
| 4d | Acquisition | +/399B (n=25)<br>+/shit$^{s}$ (n=26)<br>shi$^{ts}$/399B (n=25) | One-way ANOVA | F(2,73)=0.90, p=0.42 | N/A | N/A |
| 4e | Retrieval | +/2B (n=19)<br>+/shit$^{s}$ (n=19)<br>shi$^{ts}$/2B (n=19) | One-way ANOVA | F(2,54)=2.05, p=0.14 | N/A | N/A |
| 4f | Retrieval | +/210B (n=29)<br>+/shit$^{s}$ (n=29)<br>shi$^{ts}$/210B (n=26) | One-way ANOVA | F(2,81)=0.52, p=0.60 | N/A | N/A |
| 4g | Retrieval | +/11B (n=19)<br>+/shit$^{s}$ (n=19)<br>shi$^{ts}$/11B (n=18) | One-way ANOVA | F(2,53)=0.40, p=0.67 | N/A | N/A |
| 4h | Retrieval | +/399B (n=22)<br>+/shit$^{s}$ (n=19)<br>shi$^{ts}$/399B (n=21) | One-way ANOVA | F(2,59)=5.62, p=0.006 | Tukey | +/ shi$^{ts}$ vs +/399B p=0.93<br>+/ shi$^{ts}$ vs shi$^{ts}$/399B p=0.010<br>+/399B vs shi$^{ts}$/399B p=0.02 |
| 4j | dD1R1 | +/399B (n=20)<br>+/dD1R1(n=18)<br>dD1R1/399B (n=21) | One-way ANOVA | F(2,55)=1.767, p=0.18 | N/A | N/A |

| | | | | | | |
|---|---|---|---|---|---|---|
| 4k | dD1R1;dD1R2 | +/399B (n=14)<br>+/dD1R1;dD1R2 (n=15)<br>dD1R1;dD1R2/399B (n=15) | One-way ANOVA | F(2,41)=0.223, p=0.801 | N/A | N/A |
| 4l | dD2R | +/399B (n=23)<br>+/dD2R (n=23)<br>dD2R/399B (n=23) | One-way ANOVA | F(2,65)=0.032, p=0.968 | N/A | N/A |
| 5b | dD1R1i;dD1R2i | +/74C (n=26)<br>+/dD1R1;dD1R2 (n=28)<br>dD1R1;dD1R2/74C (n=28) | One- way ANOVA | F(2,79)=0.123, p=0.884 | N/A | N/A |
| 5b | dD2R | +/74C (n=26)<br>+/dD2Ri (n=23)<br>dD2Ri/74C (n=25) | One-way ANOVA | F(2,71)=3.51, p=0.04 | Tukey | +/ D2Ri vs +/74C p=0.47<br>+/ D2Ri vs D2Ri/74C p=0.03<br>+/74C vs D2Ri/74C p=0.29 |
| 5C | Acquisition | +/74C (n=11)<br>+/$shit^{s}$ (n=12)<br>$shi^{ts}$/74C (n=12) | One-way ANOVA | F(2,32)=0.30, p=0.75 | Tukey | N/A |
| 5D | Retrieval | +/74C (n=32)<br>+/$shit^{s}$ (n=30)<br>$shi^{ts}$/74C (n=32) | One-way ANOVA | F(2,91)=2.22, p=0.11 | Tukey | N/A |
| 5E | Consolidation | +/74C (n=21)<br>+/$shit^{s}$ (n=21)<br>$shi^{ts}$/74C (n=21) | One-way ANOVA | F(2,71)= 3.51, p=0.04 | Tukey | +/ $shi^{ts}$ vs +/74C p=0.46<br>+/ $shi^{ts}$ vs $shi^{ts}$/74C p=0.14<br>+/74C vs $shi^{ts}$/74C p=0.008 |
| 6b | Acquisition | +/18B (n=20)<br>+/$shit^{s}$ (n=21)<br>$shi^{ts}$/18B (n=25) | One-way ANOVA | F(2,63)=2.18, p=0.12 | N/A | N/A |
| 6c | Retrieval | +/18B (n=36)<br>+/$shit^{s}$ (n=38)<br>$shi^{ts}$/18B (n=45) | One-way ANOVA | F(2,116)=19.46, p=5.17x10^-08 | Tukey | +/ $shi^{ts}$ vs +/18B p=0.40<br>+/ $shi^{ts}$ vs $shi^{ts}$/18B p=0.00004<br>+/18B vs $shi^{ts}$/18B p=0.0000001 |
| 6d | Acquisition | +/58B (n=28)<br>+/$shit^{s}$ (n=30)<br>$shi^{ts}$/58B (n=30) | One-way ANOVA | F(2,85)=0.202, p=0.817 | N/A | N/A |
| 6e | Retrieval | +/58B (n=18)<br>+/$shit^{s}$ (n=20)<br>$shi^{ts}$/58B (n=22) | One-way ANOVA | F(2,57)=3.612, p=0.03 | Tukey | +/ $shi^{ts}$ vs +/58B p=0.68<br>+/ $shi^{ts}$ vs $shi^{ts}$/58B p=0.18<br>+/58B vs $shi^{ts}$/58B p=0.03 |
| 6h | dD2R | +/18B (n=20)<br>+/D2Ri (n=20)<br>D2Ri/18B (n=20) | One-way ANOVA | F(2,57)=0.113, p=0.90 | N/A | N/A |
| 6i | dD1R1;dD1R2 | +/18B (n=14)<br>+/D1R1i;D1R2i (n=14)<br>D1R1iD1R2i/18B (n=12) | One-way ANOVA | F(2,37)=2.00, p=0.15 | N/A | N/A |
| S.2 | Dopamine Acquisition and Retrieval | $shi^{ts}$/042B (n=11)<br>$shi^{ts}$/196B (n=11)<br>$shi^{ts}$/299B (n=6) | Kruskal-Wallis | $\chi^2(18)$=30.81, p=0.03 | Dunnett's Test | $shi^{ts}$/042B vs $shi^{ts}$/PBP p=0.38<br>$shi^{ts}$/196B vs $shi^{ts}$/PBP p=0.64<br>$shi^{ts}$/299B vs $shi^{ts}$/PBP p=0.55 |

| | | | | | | |
|---|---|---|---|---|---|---|
| | | shi$^{ts}$/047B (n=11)<br>shi$^{ts}$/195B (n=12)<br>shi$^{ts}$/316B (n=12)<br>shi$^{ts}$/312B (n=10)<br>shi$^{ts}$/194B (n=12)<br>shi$^{ts}$/025B (n=11)<br>shi$^{ts}$/043B (n=11)<br>shi$^{ts}$/213B (n=11)<br>shi$^{ts}$/301B (n=12)<br>shi$^{ts}$/040B (n=11)<br>shi$^{ts}$/087C (n=12)<br>shi$^{ts}$/315C (n=12)<br>shi$^{ts}$/109B (n=12)<br>shi$^{ts}$/188B (n=10)<br>shi$^{ts}$/032B (n=11)<br>shi$^{ts}$/PBP (n=11) | | | | shi$^{ts}$/047B vs shi$^{ts}$/PBP p=0.71<br>shi$^{ts}$/195B vs shi$^{ts}$/PBP p=0.79<br>shi$^{ts}$/316B vs shi$^{ts}$/PBP p=0.71<br>shi$^{ts}$/312B vs shi$^{ts}$/PBP p=0.49<br>shi$^{ts}$/194B vs shi$^{ts}$/PBP p=0.71<br>shi$^{ts}$/025B vs shi$^{ts}$/PBP p=0.17<br>shi$^{ts}$/043B vs shi$^{ts}$/PBP p=0.86<br>shi$^{ts}$/213B vs shi$^{ts}$/PBP p=0.67<br>shi$^{ts}$/301B vs shi$^{ts}$/PBP p=0.27<br>shi$^{ts}$/040B vs shi$^{ts}$/PBP p=0.54<br>shi$^{ts}$/087C vs shi$^{ts}$/PBP p=0.17<br>shi$^{ts}$/315C vs shi$^{ts}$/PBP p=0.04<br>shi$^{ts}$/109B vs shi$^{ts}$/PBP p=0.03<br>shi$^{ts}$/188B vs shi$^{ts}$/PBP p=0.008<br>shi$^{ts}$/032B vs shi$^{ts}$/PBP p=0.01 |
| S.2 | MB Acquisition and Retrieval | shi$^{ts}$/010B (n=10)<br>shi$^{ts}$/152B (n=12)<br>shi$^{ts}$/364B (n=11)<br>shi$^{ts}$/009B (n=12)<br>shi$^{ts}$/417B (n=11)<br>shi$^{ts}$/005B (n=12)<br>shi$^{ts}$/370B (n=12)<br>shi$^{ts}$/461B (n=12)<br>shi$^{ts}$/008B (n=11)<br>shi$^{ts}$/371B (n=12)<br>shi$^{ts}$/PBP (n=12) | Kruskal-Wallis | $\chi^2$(10)=27.97, p=0.002 | Dunnett's Test | shi$^{ts}$/010B vs shi$^{ts}$/PBP p=0.04<br>shi$^{ts}$/152B vs shi$^{ts}$/PBP p=3.69x10^-05<br>shi$^{ts}$/364B vs shi$^{ts}$/PBP p=0.04<br>shi$^{ts}$/009B vs shi$^{ts}$/PBP p=4.86x10^-04<br>shi$^{ts}$/417B vs shi$^{ts}$/PBP p=2.68x10^-04<br>shi$^{ts}$/005B vs shi$^{ts}$/PBP p=0.002<br>shi$^{ts}$/370B vs shi$^{ts}$/PBP p=0.04<br>shi$^{ts}$/461B vs shi$^{ts}$/PBP p=0.04<br>shi$^{ts}$/008B vs shi$^{ts}$/PBP p=1.87x10^-04<br>shi$^{ts}$/371B vs shi$^{ts}$/PBP p=0.045 |
| S.3A | Acquisition | +/HL9 (n=8)<br>+/shit$^{s}$ (n=8)<br>Shi$^{ts}$/HL9 (n=8) | One-way ANOVA | F(2,21)=0.24, p=0.788 | N/A | N/A |
| S.3B | Consolidation | +/HL9 (n=8)<br>+/shit$^{s}$ (n=8)<br>Shi$^{ts}$/HL9 (n=8) | One-way ANOVA | F(2,21)=0.698, p=0.509 | N/A | N/A |
| S.3C | Retrieval | +/HL9 (n=8)<br>+/shit$^{s}$ (n=8)<br>Shi$^{ts}$/HL9 (n=8) | One-way ANOVA | F(2,21)=8.596, p=0.00187 | Tukey | +/ shi$^{ts}$ vs +/HL9 p=0.92<br>+/ shi$^{ts}$ vs shi$^{ts}$/HL9 p=0.003<br>+/HL9 vs shi$^{ts}$/HL9 p=0.007 |
| S.4 | Dopamine Fluorescence | Air (n=7)<br>Ethanol (n=11) | One-way ANOVA | F(1,16)=2.947, p=0.105 | N/A | N/A |
| S.5 | dDop1R1 | RNAi/Elav (n=6)<br>RNAi/+ (n=6) | One-way ANOVA | F(1,10)=20.05, p=0.001 | N/A | N/A |
| S.5 | dDop1R2 | RNAi/Elav (n=6)<br>RNAi/+ (n=6) | One-way ANOVA | F(1,10)=30.31, p=0.0002 | N/A | N/A |
| S.5 | dD2R | RNAi/Elav (n=3)<br>RNAi/+ (n=3) | One-way ANOVA | F(1,4)=19.14, p=0.01 | N/A | N/A |
| S.6A | R58E02 Temperature Controls | +/R58E02 (n=25)<br>+/shit$^{s}$ (n=24)<br>Shi$^{ts}$/R58E02 (n=25) | One-way ANOVA | F(2,42)=1.953, p=0.155 | N/A | N/A |
| S.6B | HL9 Temperature Controls | +/HL9 (n=8)<br>+/shit$^{s}$ (n=8)<br>Shi$^{ts}$/HL9 (n=8) | One-way ANOVA | F(2,21)=0.823, p=0.453 | N/A | N/A |
| S.6C | 109B Temperature Controls | +/109B (n=35)<br>+/shit$^{s}$ (n=35)<br>shi$^{ts}$/109B (n=35) | One-way ANOVA | F(2,109)=0.411, p=0.664) | N/A | N/A |
| S.6D | 58B Temperature Controls | +/58B (n=18)<br>+/shit$^{s}$ (n=18)<br>shi$^{ts}$/58B (n=18) | One-way ANOVA | F(2,50)=0.516, p=0.6 | N/A | N/A |

| | | | | | | |
|---|---|---|---|---|---|---|
| S.6E | 18B Temperature Controls | +/18B (n=24)<br>+/shit$^{s}$ (n=25)<br>shi$^{ts}$/18B (n=25) | One-way ANOVA | F(2,71)=0.225, p=0.799 | N/A | N/A |
| S.6F | 399B Temperature Controls | +/399B (n=18)<br>+/shit$^{s}$ (n=18)<br>shi$^{ts}$/399B (n=18) | One-way ANOVA | F(2,51)=1.039, p=0.361 | N/A | N/A |

Supplementary Table 3. Statistical Analysis Summary for Figures 1-7 and Supplemental Figure 2-6

| Oligonucleotide Name | Sequence |
|---|---|
| CG13646F | AGTTTGACATCCACCCCGTC |
| CG13646R | CTCACTGGCGATTCCGATGA |
| Dop2RF | CTGAACTGCACCAACGAGACGC |
| Dop2RR | CAGGATGTTGCCGAAGAGGGTC |
| Dop1R1F | CCGTCGTGTCCAGCTGTATCAG |
| Dop1R1R | CTTCTCGGCCACCTCACCTG |
| Dop1R2F | CCTGGCTCGGCTGGATCAAC |
| Dop1R2R | ATCGTGGGCTGGTACTTGCG |

Supplementary Table 4. Primers for RT-qPCR

| Target Gene | Stock # | Citation |
|---|---|---|
| *Dop1R1* | VDRC-KK-107058 | (Wang, Pu et al. 2013, Agrawal and Hasan 2015, Wang, Lin et al. 2016, Ferguson, Petty et al. 2017, Lark, Kitamoto et al. 2017) |
| *Dop1R2* | VDRC-GD-3391 | (Dietzl, Chen et al. 2007, Regna, Kurshan et al. 2016, Wang, Lin et al. 2016) |
| *D2R* | VDRC-GD-11471 | (Neckameyer and White 1993, Dietzl, Chen et al. 2007, Bang, Hyun et al. 2011, Shang, Haynes et al. 2011, Agrawal and Hasan 2015, Wang, Lin et al. 2016, Andreatta, Kyriacou et al. 2018, Petruccelli, Feyder et al. 2018) |

Supplementary Table 5. Previous publications using RNAi lines in this paper.
